# The Single Cell Proteomic blueprint, navigating instrumentation platforms, software tools and high-load libraries in neutrophils, RKO and A549 cells

**DOI:** 10.64898/2026.06.12.731618

**Authors:** Alejandro J Brenes, Rupert L Mayer, Agata Makar, Patricia Coelho, Gabi van Stralen, Pranvera Sadiku, Sarah R Walmsley, Manuel Matzinger, Karl Mechtler, Alex von Kriegsheim

## Abstract

Mass spectrometry-based single cell proteomics (SCP) is rapidly emerging as a powerful approach for biological research, with applications extending beyond in-vitro cancer cell lines. Recent advances make it possible to apply SCP to ex-vivo human cells from tissues such as the brain and pancreas, as well as to technically challenging immune populations such as neutrophils. However, these analyses remain more challenging and typically result in reduced proteomic coverage. To support the development of robust workflows for SCP data acquisition and analysis, we systematically evaluated multiple DIA search engines, search engine settings, the inclusion of high-load library samples in single-cell search spaces, the impact of contaminants, and the quantitative properties of identified proteins. These comparisons were performed across two major instrumentation platforms, Orbitrap Astral and timsTOF SCP, and across A549, RKO cells and neutrophils, three cell types differing in size and protein content. Our work here provides guidelines on the software parameters to use for SCP, instrument specific results and cell dependent optimizations of high-load libraries, as well as novel evaluation of the quantitative properties of proteins for single cell and low input proteomics.

## Introduction

Mass spectrometry-based single cell proteomics (SCP) is a very powerful technique to explore cell populations, functional states and phenotypes^1,2^. The first pioneering implementations used data-dependent acquisition (DDA) methods paired with tandem mass tag (TMT) labelling^3,4^. However, there are significant limitations to DDA in general and TMT in particular^5^. Due to improvements in software solutions to interpret data independent acquisition (DIA) data^6,7^, label-free DIA-based approaches have become the more common and robust workflows^8-14^.

Due to a lack of amplification steps in SCP, the earlier data was limited mostly to high protein content in-vitro cell lines^9,11^. However, with recent improvements in instrument sensitivity^11,12,15^, sample processing^8,16-19^ as well as software tools^20^, it has recently become possible to study primary human cells, ranging from brain cells^21^, to pancreatic islets^22,23^, hematopoietic stem cells^12,24^ to even challenging immune cells like neutrophils^25^.

As SCP expands to more novel biological applications, it is important to consider how to best design SCP experiments that improve data quality, proteomic coverage and biological interpretability. To develop improved workflows for SCP data acquisition and analysis, we compared the main DIA search engines, DIA-NN and Spectronaut, evaluated different software settings, and assessed the impact of including high-load library samples within the single-cell search space. These analyses were performed across the two main SCP instrumentation platforms: Orbitrap Astral (Astral) and timsTOF SCP (tSCP).

These analyses were carried out across three distinct types of cells of varying diameter and total protein content (Fig. 1A). We first studied A549 cells, a lung adenocarcinoma-like cell line with an estimated protein content between 200-250pg and a diameter range of 10-30µm^26^. These cells are a typical use case representing larger human cells with higher protein content. We also analysed RKO cells, colorectal carcinoma cell line with an estimated 150-200pg of protein content and 14-20µm diameter^27^. As medium-sized cells, RKO cells are more challenging to analyse than A549 cells. Finally, we analysed peripheral blood human neutrophils, which have an estimated protein content of 30-60 pg and a diameter of 8-10 µm^28^ (Fig. 1A). Neutrophils are among the most challenging human cells to study by SCP because of the low protein content, restricted proteome and high dynamic range^1,25,29^. Together, these cell types cover key scenarios that represent most current SCP use cases.

**Figure 1.**
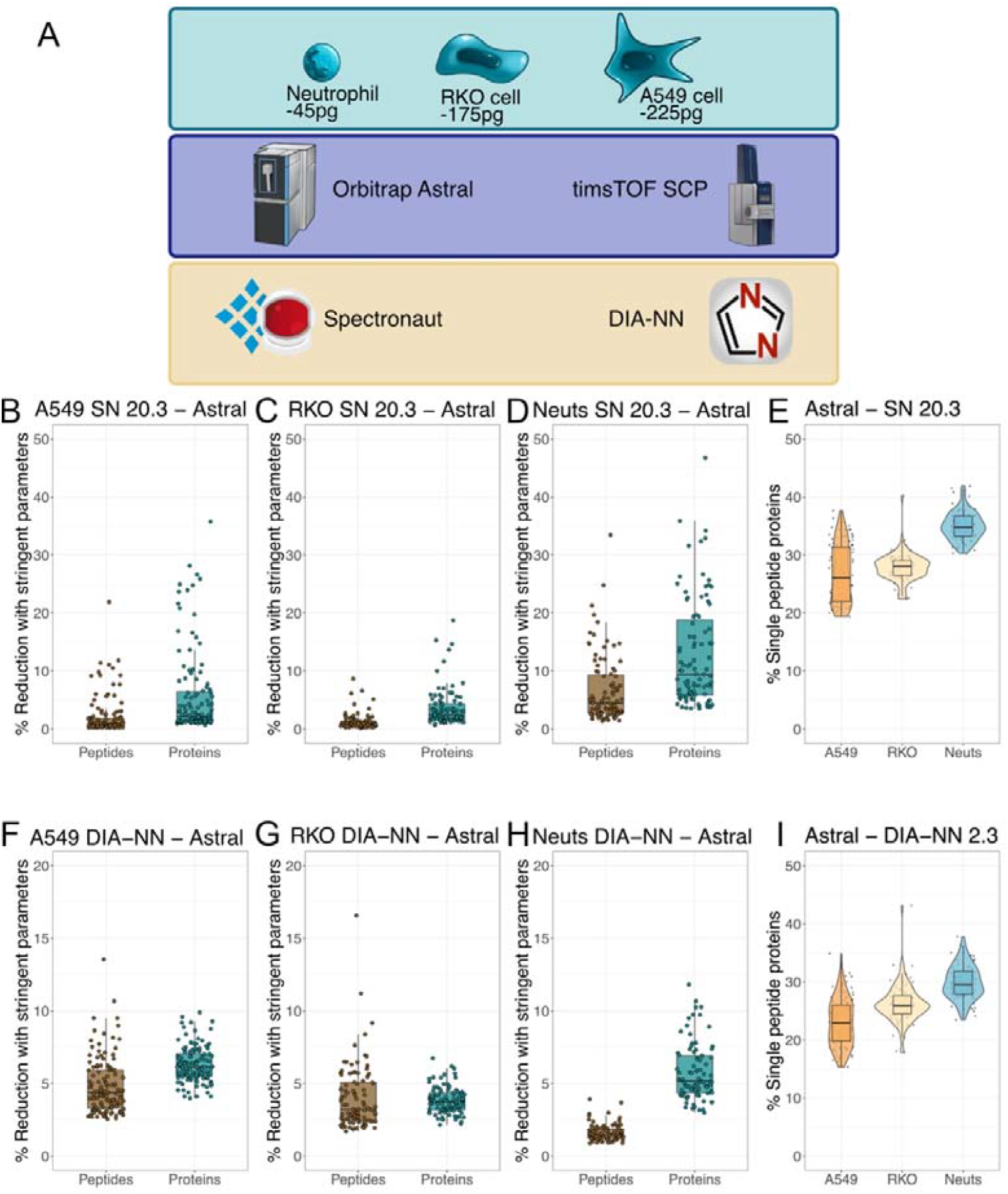
Identification rates across parameters settings for the Astral: **(A)** Schematic showing the cell types, instruments and softwares used. Boxplots showing the number of percentage reduction in peptide and protein identifications when using the stringent settings vs the default in Spectronaut (SN) across **(B)** A549 **(C)** RKO cells **(D)** Neutrophils (Neuts). **(E)** Boxplot showing the percentage of proteins identified by a single peptide with SN across all 3 cell types. Boxplots showing the number of percentage reduction in peptide and protein identifications when using the stringent settings vs the default in DIA-NN across **(F)** A549 cells, **(G)** RKO cells, **(H)** Neuts. **(I)** Boxplot showing the percentage of proteins identified by a single peptide with DIA-NN across all 3 cell types. For all boxplots, the top and bottom hinges represent the 1st and 3rd quartiles. The top whisker extends from the hinge to the largest value no further than 1.5× interquartile range (IQR) from the hinge; the bottom whisker extends from the hinge to the smallest value at most 1.5× IQR of the hinge

Across these cell types, we compared protein identification rates, quantification precision and linearity, the effects of including high-load library samples in the search space as well as the impact of contaminants. Our data provides a comprehensive evaluation of instrumentation platform and software tools, with optimised suggestions for the tSCP and Astral experiments. In addition, we define an extended panel of contaminant proteins that can affect quantitation and the interpretation of SCP results.

## Results

### Stringent search parameters only cause minor reductions in protein IDs for single cell

We have previously shown that setting stringent search parameters is vital to reducing false-positive identifications and improving DIA data interpretation^30^. Hence after excluding contaminants, we first tested the effects of the more stringent SCP parameters developed in our previous work^25^. The Spectronaut results show that these stringent parameters result in a median reduction in identifications of <1% at the peptide level, and 2.2% at the protein group level, referred to henceforth as protein level, in the larger A549 cells (Fig. 1B). Similarly, in RKO cells, we saw a reduction of <1% in peptide identifications and only 2.4% at the protein level (Fig. 1C). It was only in the smaller neutrophils that we observed larger effects, with a 4.5% reduction at the peptide level and ∼9% at the protein level (Fig. 1D). This was directly correlated with the percentage of single peptide proteins identified, which was highest in the neutrophil data (Fig. 1E). This shows the parameters have stronger effects in smaller cell types.

We further tested the reduction in identifications using DIA-NN. Here we found a ∼4% reduction at the peptide and ∼6% reduction at the protein level in A549 cells (Fig. 1F). RKO cells displayed ∼3% reduction of identifications at the peptide level and ∼4% at the protein level (Fig. 1G). Strikingly, unlike Spectronaut, neutrophil data from DIA-NN did not display the most pronounced effect, with only a ∼1.5% reduction at the peptide level, which was amplified to 5% at the protein level (Fig. 1H). The increased reduction at the protein level compared to the peptide level reflects the fact that neutrophils also have a higher proportion of single-peptide protein identifications (Fig. 1I). It is worth noting that the percentage of single peptide proteins in neutrophils with DIA-NN is considerably lower than in Spectronaut. This explains why DIA-NN had more moderate reductions in the protein identifications with the stringent parameters and why in DIA-NN smaller cells suffer lower effects from the more stringent parameters.

In summary, both the Spectronaut and DIA-NN data confirm that using more stringent parameters for the Astral have limited effect on the proteomic depth and should thus be used routinely for single-cell proteomics. All posterior analyses shown here use these stringent parameters.

### The Astral has better sensitivity for low protein content cells

We next focused on analysing the protein identification rates across the different cell types across both Spectronaut and DIA-NN. These were analysed both on the Astral and the tSCP. Starting with the Astral analysed in Spectronaut, we detected a median of 4,349 proteins per single A549 cell, representing a high proteomic coverage for a single cell. The detection rates follow the expected patterns across cell sizes: the medium-sized RKO cells showed lower identification rates, with a median of 3,383 per cell, while the smaller neutrophils yielded only 1,544 proteins per cell (Fig.2A). Reassuringly, the same trend was observed when the data were searched with DIA-NN, with the highest number of protein identifications in the A549, followed by a progressive reduction in identifications, with the lowest rates seen in neutrophils (Fig. 2B).

**Figure 2.**
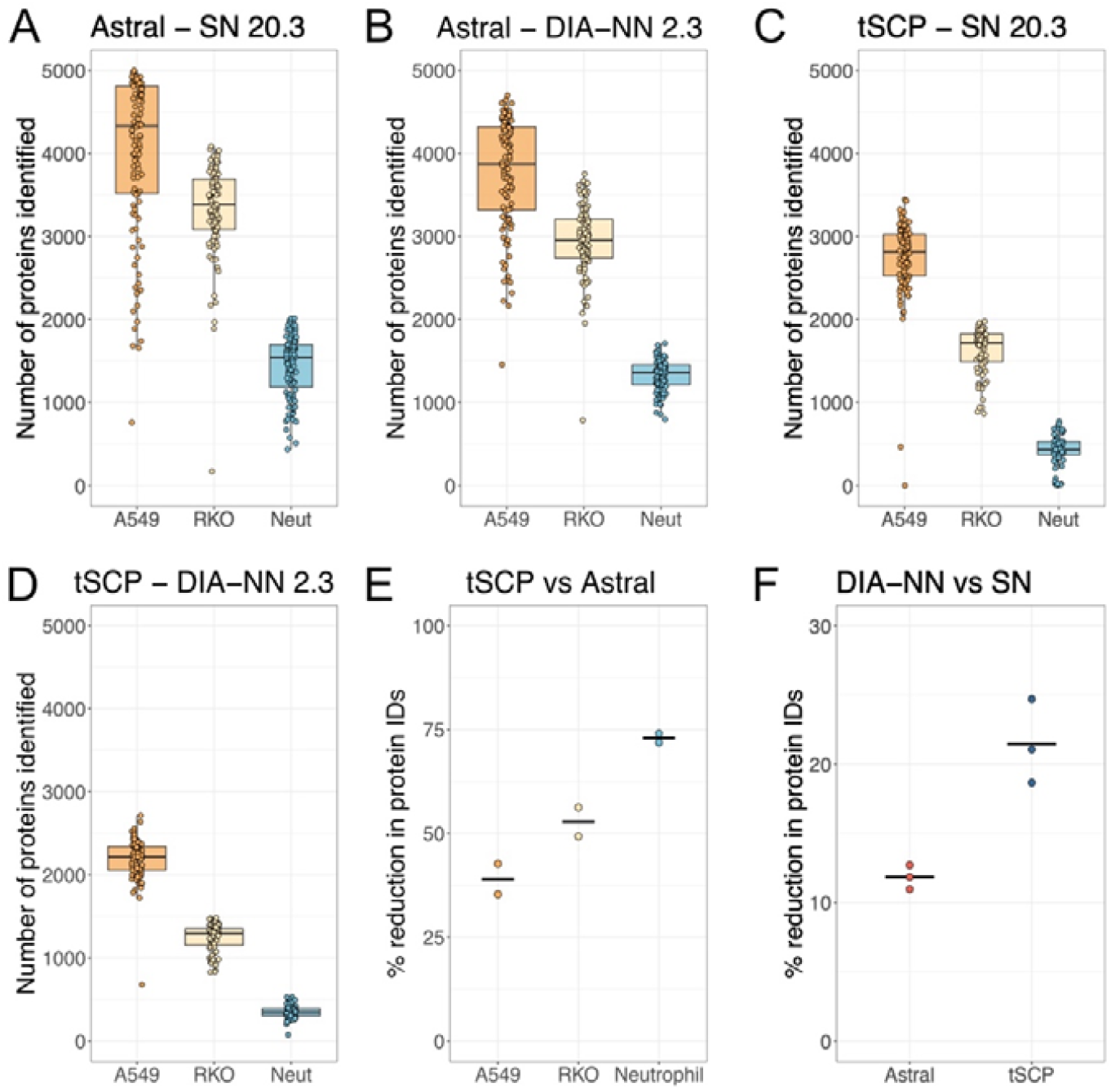
Single cell identification rates across software and instruments: (A-D) Boxplots showing the number of protein IDs over all three cell types for the Astral when using (A) Spectronaut (SN) and (B) DIA-NN. For the timTOF SCP when using (C) SN and (D) DIA-NN. (E) Graph illustrating % reductions in protein IDs when comparing the tSCP with the Astral for each cell type the two data points represent the results on SN and DIA-NN. (F) Panel visualizing the reduced number of protein IDs of DIA-NN over SN, the three data points represent the 3 cell types. For all boxplots, the top and bottom hinges represent the 1st and 3rd quartiles. The top whisker extends from the hinge to the largest value no further than 1.5× interquartile range (IQR) from the hinge; the bottom whisker extends from the hinge to the smallest value at most 1.5× IQR of the hinge

We also analysed the same three cell types on a different instrumentation platform, in this case using Trapped Ion Mobility Spectrometry (tims) on the tSCP. With Spectronaut the medium number of proteins identified in A549 cells was 2,811 proteins with clear reductions in protein identifications as the cells got smaller (Fig. 2C). The same pattern was observed with DIA-NN, showing 2,219 proteins identified on A549 cells and the lowest number of identifications in neutrophils with <500 proteins identified (Fig. 2D).

We proceeded to directly compare the two instrumentation platforms; however, it should be noted that the tSCP is an older generation instrument and newer versions have been released which have significant upgrades resulting in better sensitivity and dynamic range. The reduced sensitivity of the tSCP is reflected by the fact that the bigger cells, in this case the A549, only display a median 39% reduction in protein identifications across both software tools when compared the Astral. However, as the cells get smaller, the differences between instruments gets bigger, with the medium sized RKO cells showing a 53% reduction in protein identifications and with the much smaller neutrophils this figure inflates to 73% (Fig. 2E). suggesting that the Astral is far superior for identification rates in low protein content cells. This suggests that the Astral provides substantially higher identification rates in cell with low protein content. Therefore, for primary cells with limited protein input, the use of newer-generation instrumentation may offer significant benefits.

Finally, we also compared whether there were any systematic differences when using the two software tools across the two instrumentation platforms. Searching the Astral data with DIA-NN produced a median reduction in protein identification rates of 12%. However, we noted that this figure was significantly higher for the tSCP data, with a median decrease of 21% across all 3 cell types (Fig. 2F), suggesting Spectronaut is better optimised for timsTOF data.

### Comparing the quantification precision across instruments and software tools

After comparing identification rates across different cell types and instruments, we looked at the precision by analysing the coefficient of variation (CV) following the recommended guidelines^31^. The searches in Spectronaut were done without method evaluation and the searches in DIA-NN had match-between-runs (MBR) enabled. Without applying normalisation, we looked at the CVs of the raw intensities for all cell types across instruments and software tools, showing the real spread of the data. As A549 and RKO cells are larger and grown in-vitro we expected their CVs to be lower. The data confirmed this theory, as on the Astral using Spectronaut A549 cells showed a median CV of 55% and RKO cells 59%. The smaller ex-vivo neutrophils showed a much higher median of 114% CV (Fig. 3A). The pattern was similar when we studied the CVs produced with DIA-NN (Fig. 3B). However, in every case, DIA-NN produced significantly lower CVs compared to Spectronaut, with 11% lower CVs in A549 cells, 7% lower in RKO cells and a marked 13% reduction in CVs in neutrophils (Fig. 3C).

**Figure 3.**
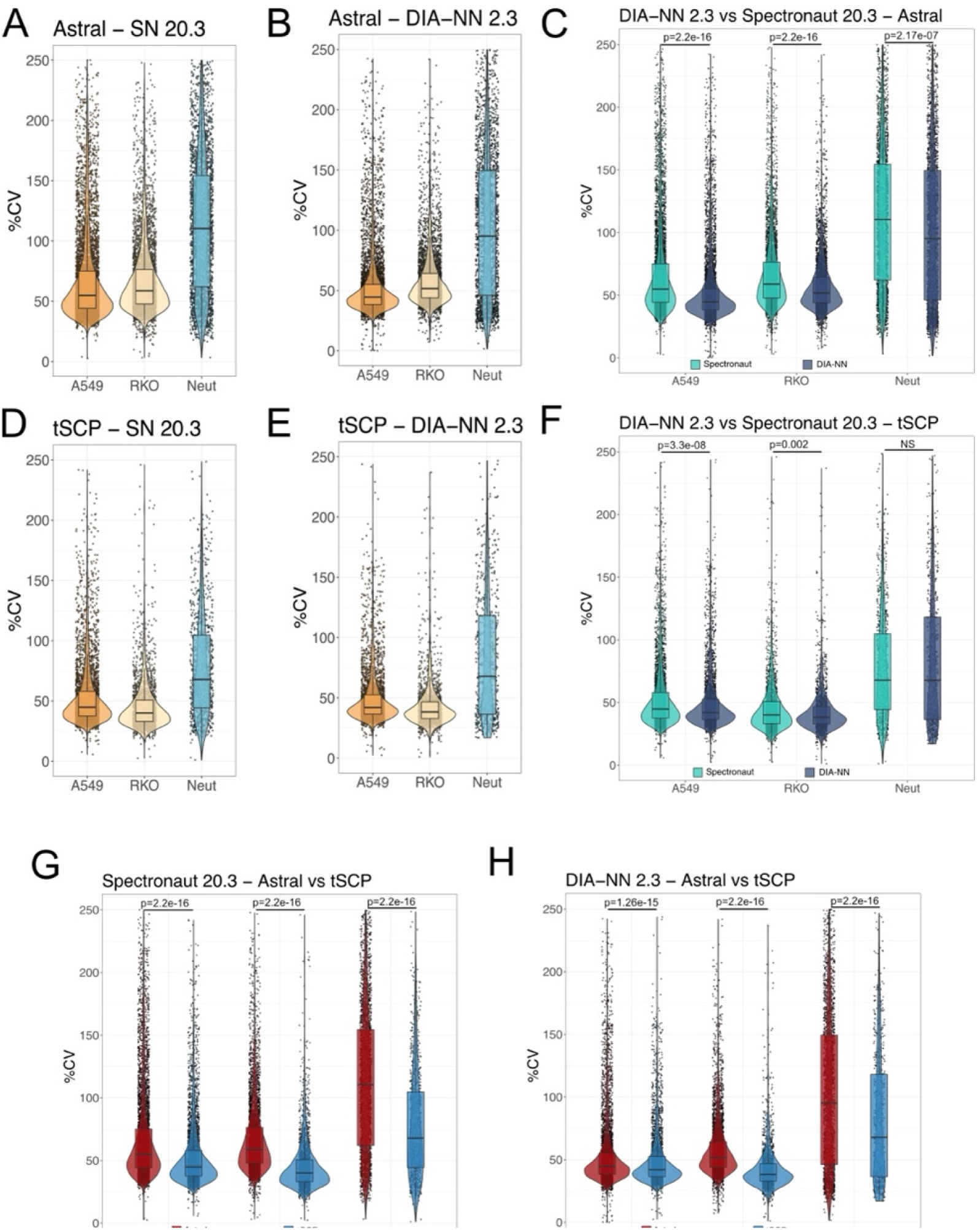
Coefficient of variation (CV) across software and instrument: Box plots showing the CV across all proteins for all single cells in: **(A)** A549 cells, RKO cells and neutrophils analysed on an Orbitrap Astral (Astral) with Spectronaut (SN), **(B)** A549 cells, RKO cells and neutrophils analysed on an Astral with DIA-NN, **(C)** direct comparison of Spectronaut and DIA-NN for an Astral across all cell types, **(D)** A549 cells, RKO cells and neutrophils analysed on a timsTOF SCP (tSCP) with Spectronaut (SN), **(E)** A549 cells, RKO cells and neutrophils analysed on a tSCP with DIA-NN, **(F)** direct comparison of Spectronaut and DIA-NN for an Astral across all cell types, **(G)** Spectronaut based Astral and tSCP comparison across cell types with matched number of proteins for both instruments, and (H) DIA-NN based Astral and tSCP comparison across cell types with matched number of proteins for both instruments. For all boxplots, the top and bottom hinges represent the 1st and 3rd quartiles. The top whisker extends from the hinge to the largest value no further than 1.5× interquartile range (IQR) from the hinge; the bottom whisker extends from the hinge to the smallest value at most 1.5× IQR of the hinge. Boxplot y axis limited to 250 for visualisation purposes.

On the tSCP a similar pattern was displayed. When using Spectronaut A549 cells had a median CV of 45%, RKO cells 40% and neutrophils a much higher 69% CV (Fig. 3D). The displayed pattern was identical when analysing the data with DIA-NN (Fig. 3E). However, for the tSCP DIA-NN produced only a marginal reduction of ∼2% CVs for both in-vitro cell types and with the CVs not being significantly different in neutrophils (Fig. 3F). This suggests that potentially the parallel accumulation and serial fragmentation capacities of the tSCP reduce interfering ions potentially, limiting the improvements seen by using QuantUMS^32^ on DIA-NN.

Finally, we directly compared the CVs between the two instrumentation platforms. Because the Astral identified a larger number of lower-abundance proteins, which are reported to have higher variability^33,34^, we restricted the Astral dataset to the most abundant proteins, so that the same number of proteins were used to calculate the CVs across both instruments. Despite this adjustment, the tSCP produced significantly lower CVs across all three cell types. The smallest difference was observed in A549 cells, where the tSCP produced 11% lower CVs with Spectronaut (Fig. 3G) and 2% with DIA-NN (Fig. 3H). The greatest improvement was observed in neutrophils where, the tSCP produced a staggering 46% lower CVs with Spectronaut (Fig. 3G) and 32% lower CVs with DIA-NN (Fig. 3H). In summary, DIA-NN increases the precision of Astral data while little effect on the tSCP data. Moreover, direct platform comparison showed that tSCP produced more precise quantitative data than the Astral, particularly in smaller single cells such as neutrophils, which is highly relevant for studies focused on primary human cells.

### High-load libraries provide no significant identification gains with Astral data

Based on previous work^12,35,36^ we also investigated how adding high-load libraries to the search space affects proteomic depth and protein quantification precision. A high-load library is of a sample with higher total protein load. However, in our case, unlike previous studies^12,13,36^ our high-load libraries did not use a bulk dilution, instead were a titration with increasing numbers of cells. This ensures a sample matrix and digestion quality fitting well to the actual single cell samples which, in theory, allow for improved matching. For the smaller neutrophils and the medium-sized RKO cells we used high-load libraries of 5, 10, 25 and 50 cells and for the larger A549 cells we used 5, 11, 20 and 40 cells. Then, both on Spectronaut and DIA-NN, we searched the single cell raw files with the addition of one type of high-load library at a time.

Using Spectronaut, we found no significant differences in the number of proteins identified in A549 cells when high-load libraries were included in the search space (Fig. 4A). However, in RKO cells, the use of 50-cell high-load libraries led to a significant decrease in protein identification rates (Fig. 4B). A similar effect was observed in neutrophils, where 25- and 50-cell libraries produced significant reductions in protein identifications of more than 16% (Fig. 4C). When the same data were analysed with DIA-NN, high-load libraries did not significantly alter protein identification rates in any cell type (Fig. 4D–F). This pattern was also observed at the peptide level (Supplementary Fig. 1). Overall, high-load libraries did not improve protein identification rates for Astral data across any software or cell type, and in some cases reduced protein identifications when the input level of the library was too high.

**Figure 4.**
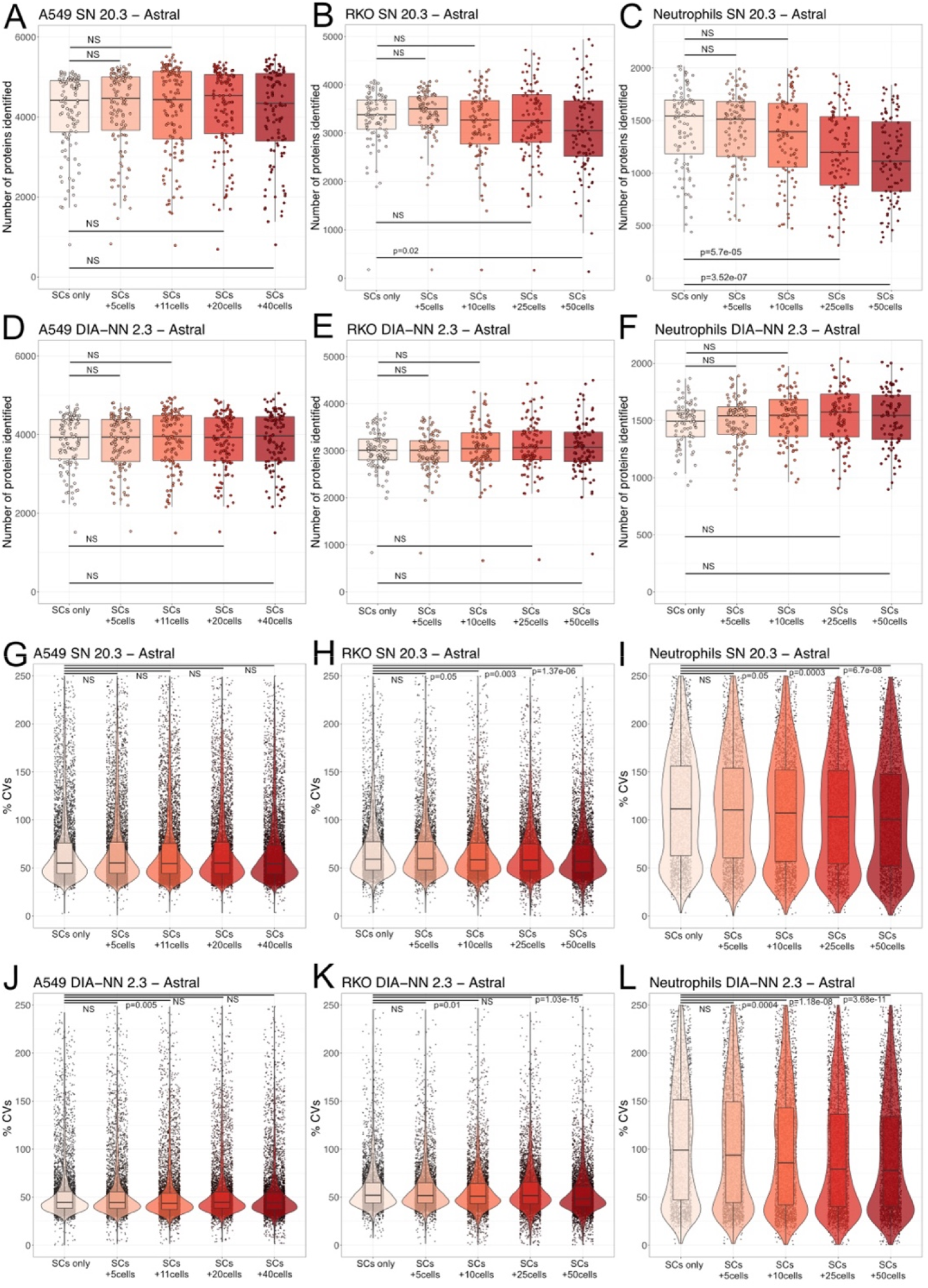
Astral IDs and CVs across high-load library search spaces: Boxplots showing the number of proteins identified on an Orbitrap Astral across the different search spaces which include single cells (SC) with the high-load libraries for **(A)** A549 cells with Spectronaut (SN), **(B)** RKO with SN, **(C)** neutrophils with SN **(D)** A549 cells with DIA-NN, **(E)** RKO cells with DIA-NN and **(F)** neutrophils with DIA-NN. Box and violin plots showing the percentage coefficient of variation (CV) across the different search spaces for **(G)** A549 cells with SN, **(H)** RKO cells with SN, **(I)** neutrophils with SN, **(J)** A549 cells with DIA-NN, **(K)** RKO cells with DIA-NN and **(L)** neutrophils with DIA-NN. For all boxplots, the top and bottom hinges represent the 1st and 3rd quartiles. The top whisker extends from the hinge to the largest value no further than 1.5× interquartile range (IQR) from the hinge; the bottom whisker extends from the hinge to the smallest value at most 1.5× IQR of the hinge. Boxplot y axis limited to 250 for visualisation purposes.

We then proceeded to analyse potential changes in the quantification, as previous studies had reported lower precision and higher dispersion in the quantification when including the high-load libraries within the search space^13^. Our data showed that for the larger A549 cells there were no significant changes in CVs when including the high-load libraries with Spectronaut (Fig. 4G). RKO cells displayed modest reductions of up to 2% in CVs using the 10, 25 and 50 cell libraries (Fig. 4H). Neutrophils displayed more considerable reductions in CVs when using the 10, 25 and 50 cell libraries with a maximum reduction ∼12% with the 50-cell library (Fig. 4I).

Similar to the Spectronaut results, the DIA-NN analysis of A549 cells showed no significant improvement in CVs when high-load libraries were used (Fig. 4J). In contrast, RKO cells showed significant changes with the 10 and 50-cell libraries (Fig. 4K). Consistent with the Spectronaut analysis, the largest improvement was observed in neutrophils, where the 10, 25, and 50-cell libraries all produced significant reductions in CVs. The greatest effect was seen with the 50-cell library, which reduced CVs by 21% (Fig. 4L).

Overall, for the Orbitrap Astral when using the appropriate more stringent search settings, high-load libraries provide no identification gains with DIA-NN 2.3 or Spectronaut 20. In Spectronaut, libraries containing more than 25 cells can reduce protein-level identification rates. In DIA-NN, high-load libraries do not significantly affect protein identifications, but they can substantially improve quantification precision in medium and small cells.

### High-load libraries improve IDs and CVs with DIA-NN on the timsTOF SCP

Similar to Astral analysis, we compared the identification rates and quantification precision on data generated from the timsTOF SCP (tSCP). As observed with the Astral data, the results for DIA-NN and Spectronaut differed. In Spectronaut, including high-load libraries of more than 10-cells in the search space significantly reduced the number of protein identifications across all cell types (Fig. 5A–C). The strongest effect was observed in neutrophils, where use of the 50-cell library reduced protein identifications by more than 44% (Fig. 5C).

**Figure 5.**
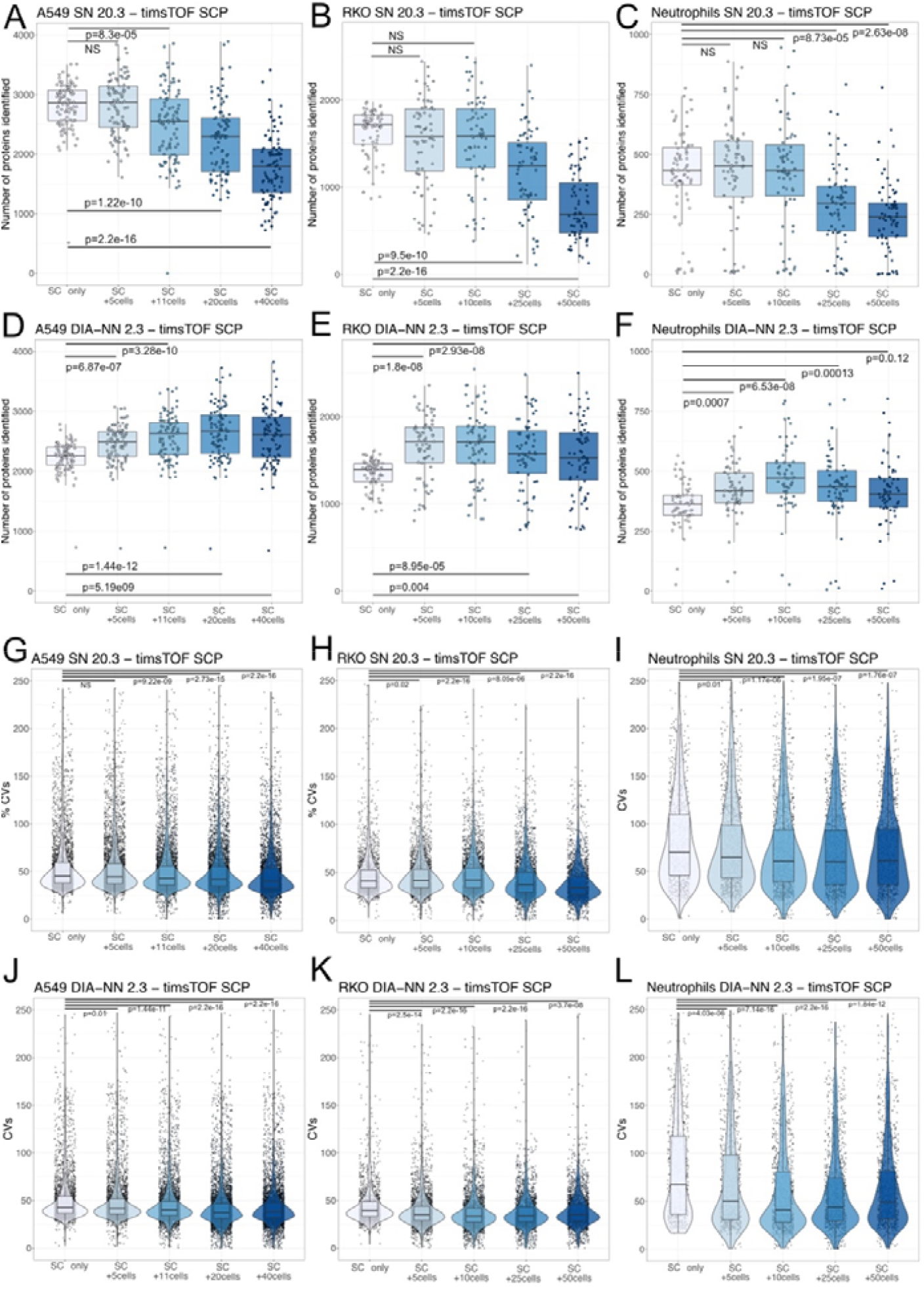
tSCP IDs and CVs across high-load library search spaces: Boxplots showing the number of proteins identified on a timsTOF SCP (tSCP) across the different search spaces which include single cells (SC) with specific high-load libraries for **(A)** A549 cells with Spectronaut, **(B)** RKO cells with Spectronaut, **(C)** neutrophils with Spectronaut, **(D)** A549 cells with DIA-NN, **(E)** RKO cells with DIA-NN and **(F)** neutrophils with DIA-NN. Box and violinplots show the percentage coefficient of variation (CV) across the different search spaces for **(G)** A549 cells with Spectronaut, **(H)** RKO cells with Spectronaut, **(I)** neutrophils with Spectronaut, **(J)** A549 cells with DIA-NN, **(K)** RKO cells with DIA-NN and **(L)** neutrophils with DIA-NN. For all boxplots, the top and bottom hinges represent the 1st and 3rd quartiles. The top whisker extends from the hinge to the largest value no further than 1.5× interquartile range (IQR) from the hinge; the bottom whisker extends from the hinge to the smallest value at most 1.5× IQR of the hinge. Boxplot y axis limited to 250 for visualisation purposes.

The opposite pattern was seen with DIA-NN, where all cell types showed a significant increase in protein identifications when high-load libraries were included in the search space (Fig. 5D-F). The 10-cell library increased identifications by 28% in RKO cells and by 24% in neutrophils, while the 20-cell library increased protein identifications by 18% in A549 cells (Fig. 5D). Thus, for tSCP data analysed with DIA-NN, including high-load libraries in the search space significantly improved protein identification rates. This trend was also observed at the peptide level (Supplementary Fig. 2).

We next studied the precision of the quantification. With Spectronaut CVs are improved across all cell types when using high-load libraries, but at the cost of reduced protein identifications. The biggest reductions in CVs were seen with the 40&50-cell libraries dropping CVs by 5% in A549 cells (Fig. 5G), 7% in RKO cells (Fig. 5H) and 9% in neutrophils (Fig. 5I). High-load libraries also reduced the CVs with DIA-NN, with the 40-cell library reducing CVs by 4% in A549 cells (Fig. 5J), the 50-cell reducing CVs by 2% in RKO cells (Fig. 5K) and the 10-cell library reducing CVs in neutrophils by 28% (Fig. 5L).

In summary, for the tSCP data, high-load libraries provide little benefit when analysed on Spectronaut. In contrast, when analysed in DIA-NN they significantly increase the number of proteins identified while also reducing CVs, making them highly advantageos. For DIA-NN 2.3 high-load library is between 10 and 20 cells.

### Astral and timsTOF SCP display comparable identification rates when injecting over 3 nanograms

With the stark differences between the Astral and tSCP at the single cell level, we test the potential differences in performance as increased the number of cells analysed. We performed new searches across all 3 cell types, including the single cell files and all the high-load libraries within a single search space and analysed them on DIA-NN and Spectronaut.

As previously mentioned, the Astral has significantly higher sensitivity, showing >50% increased identifications at the single cell level (Fig. 2E). We were interested to understand how the differences in performance fared across the different cell numbers. We used a self-starting nonlinear model (SSasymp) to fit an asymptotic growth curve to the number of proteins identified across the different cell types. For the model we protein inputs of 200pg for A549 cells, 150pg for RKO cells and 45pg for neutrophils. Our data suggested that, for both DIA-NN and Spectronaut, the protein identification rates of the Astral and the tSCP converge at around 13 A549 cells (Fig. 6A&B) which is the equivalent of ∼2.6ng. Beyond 2.6ng the identification rates of the two instruments are comparable.

**Figure 6.**
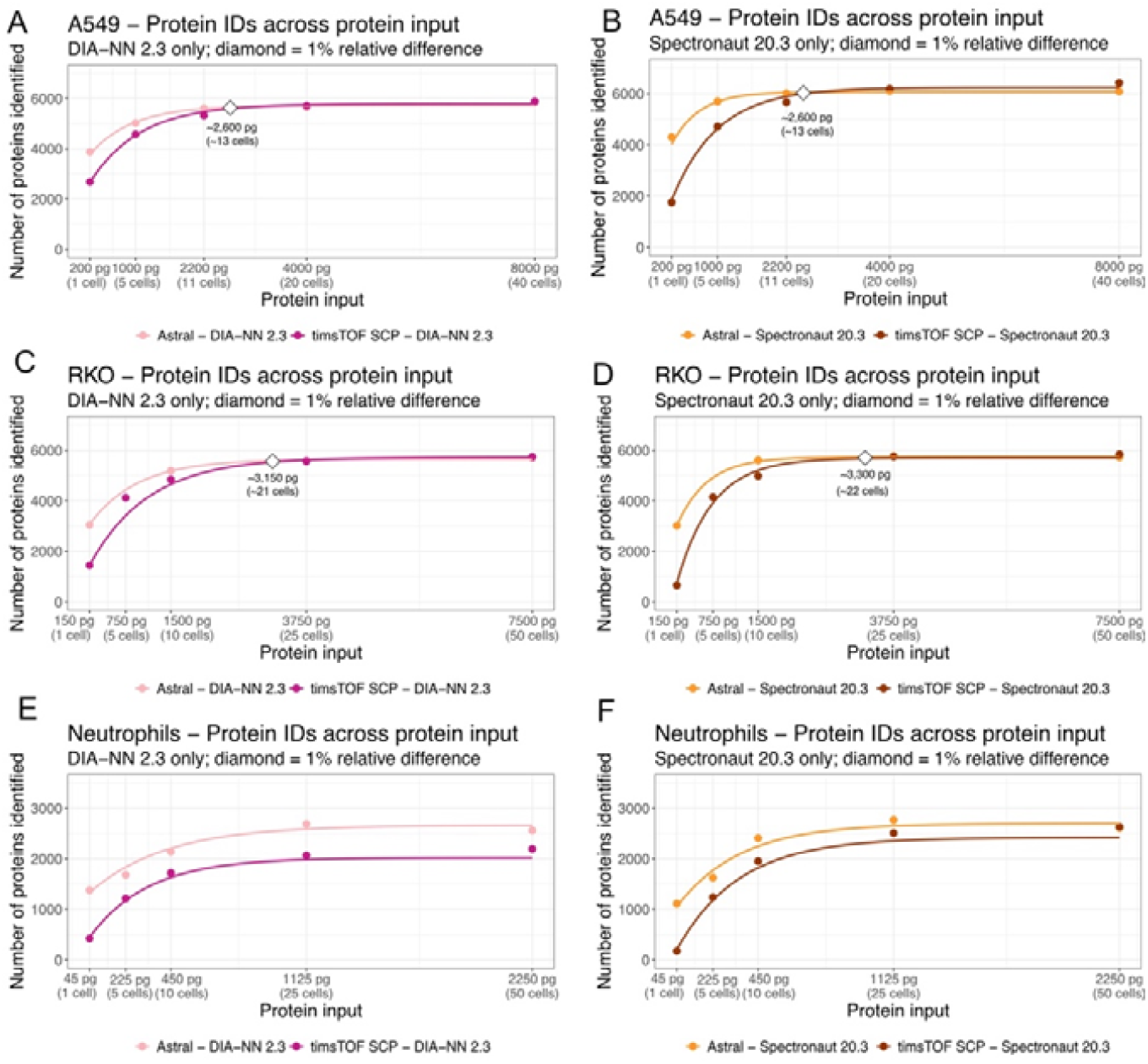
head to head instrument comparison: Line plots showing the self-starting nonlinear model (SSasymp) asymptotic growth curve showing the number of proteins identified over the estimated protein input load for **(A)** A549 cells using DIA-NN, **(B)** A549 cells using Spectronaut, **(C)** RKO cells using DIA-NN, **(D)** RKO cells using Spectronaut, **(E)** Neutrophils using DIA-NN and **(F)** neutrophils using Spectronaut. The dots show the median protein identifications across cells and loads.

We performed the same analysis on the medium-sized RKO cells. RKOs showed a consistent pattern across both Spectronaut and DIA-NN. Using DIA-NN, instrument performance converges at approximately 21 cells (Fig. 6C), and at approximately 22 cells when using Spectronaut (Fig. 6D). Thus, we estimate beyond 3.2ng the identification rates of the two instruments are comparable.

For the smaller neutrophils, no convergence was observed within this dataset, even at the 50 cells level (Fig. 6E&F). We hypothesise this is because at 50 neutrophils correspond only a total of ∼2.25ng would be injected into the mass spectrometer, which is below the predicted convergence point for both A549 and RKO cells. Based on our data, we estimate that approximately 66 neutrophils would be the point where the performance of the Astral and the tSCP might converge. In short, the Astral has significantly increased sensitivity at the single cell level, however this superior identification performance diminished with higher loads, with the tSCP displaying comparable identification rates at ∼3ng.

### Limit of quantification and the effect of contaminants for single cell and low cell numbers

We were also interested in assessing the quantitative reliability of the identified proteins and therefore applied an approach conceptually similar to the matched matrix background method^37^, except in our case we wanted to explore the linear increases in the intensity of proteins as the number of cells increased. For this analysis we focussed exclusively on DIA-NN, however the Spectronaut data is available in Supplemental Figure 3. Proteins with better quantification would display linear signatures, increasing their raw intensity as the number of cells analysed increased. To test this, we used the Jonckheere-Terpstra (JT) test which directly measures trends over ordered groups and produced a relative score. A relative JT score of 0.5 means virtually no increase in intensity as the number of cells increases, and a score 1 of almost a perfect monotonic increase.

First, we analysed the total LFQ intensity across all different cell loading conditions. On the Astral we found excellent concordance, with relative JT scores > 0.94 (Fig. 7A-C). The pattern on the tSCP was even more consistent as even neutrophils, the smallest and most challenging cells, showed a relative JT score >0.99 (Fig. 7D-F). The tSCP result shows almost perfect monotonic increase. However, like in previous analysis, we had filtered out all contaminants before analysing this data. We next wanted to evaluate the effects that contaminants could have on the quantitation. Hence, performed the analysis again without filtering the proteins contained within the contaminants database.

**Figure 7.**
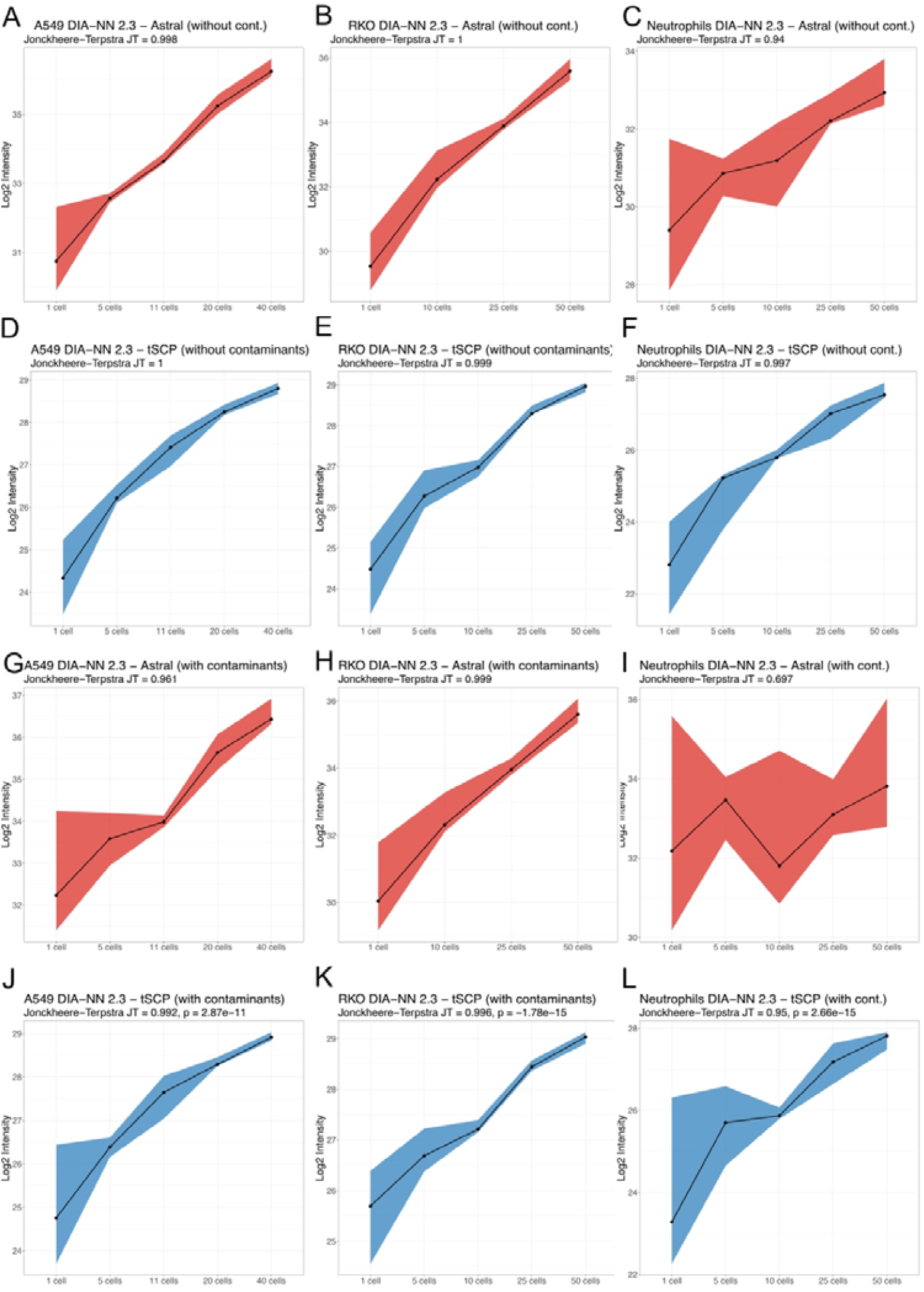
DIA-NN Intensity scaling with and without contaminants: Line plots showing the total intensity from DIA-NN across all proteins vs the different number of cells for **(A)** A549 on the Orbitrap Astral (Astral) without contaminants, **(B)** RKO on the Astral without contaminants, **(C)** Neutrophils on the Astral without contaminants, **(D)** A549 on the timsTOF SCP (tSCP) without contaminants, **(E)** RKO on the tSCP without contaminants, **(F)** neutrophils on the tSCP without contaminants, **(G)** A549 on the Astral with contaminants, **(H)** RKO on the Astral with contaminants, **(I)** Neutrophils on the Astral with contaminants, **(J)** A549 on the tSCP with contaminants, **(K)** RKO on the tSCP with contaminants, **(L)** neutrophils on the tSCP with contaminants. The dots represent the median intensity across the cell numbers and the ribbons the extent of the 95% confidence interval of the intensity.

Contaminants databases can comprise over 200 proteins, with the main components being keratins. Keratins are important components of the skin and be highly abundant contaminants even in bulk. However, keratins are also part of the epithelial proteomes. With RKO and A549 cells being of epithelial origin, and having increased protein content, it was expected contaminants might have a reduced effect. On the Astral, the data confirmed this theory, with only a reduction of 0.032 in the relative JT score for A549 cells (Fig. 7G), and virtually no effect in RKO cells (Fig. 7H). However, the effect on neutrophils was very strong, reducing the relative JT score by ∼0.24 (Fig. 7I). The effect looked far less pronounced on the timsTOF (Fig. 7J-L), where even in neutrophils the reduction was only of ∼0.05 (Fig. 7L).

In summary the data shows excellent monotonic increases in intensity as the number of cells increases, however the contaminants play an important quantitative disruption on smaller cells, in particular when analysing them on the Astral. This effect appears less prominent on the tSCP, however this might be caused by the reduced sensitivity of the instrument.

### An extended contaminant database for single cell proteomics

To further explore the effects of contaminants we focussed on doing the relative JT score of all proteins contained within the contaminants database. Because the neutrophil proteomes express no keratins, we used them as a baseline cell population. We noted that for both the Astral (Fig. 8A) and the tSCP (Fig. 8B) DIA-NN produced significantly better relative JT scores, so we focused on the DIA-NN results. We find that for both the Astral (Fig. 8C) and the tSCP (Fig. 8D) 90% of proteins on the contaminants database have a relative JT score <0.7. Suggesting contaminants have poor monotonic increases in intensity as the number of cells increase, as would be expect. Interestingly in A549 cells we could detect a high proportion of contaminant proteins with very high relative JT scores (>0.9), and this was consistent across both the Astral (Fig. 8E) and the tSCP (Fig. 8F). However most of these proteins are keratins, and there is solid evidence that they are expressed in A549 cells^38-40^, explaining the high scores.

**Figure 8.**
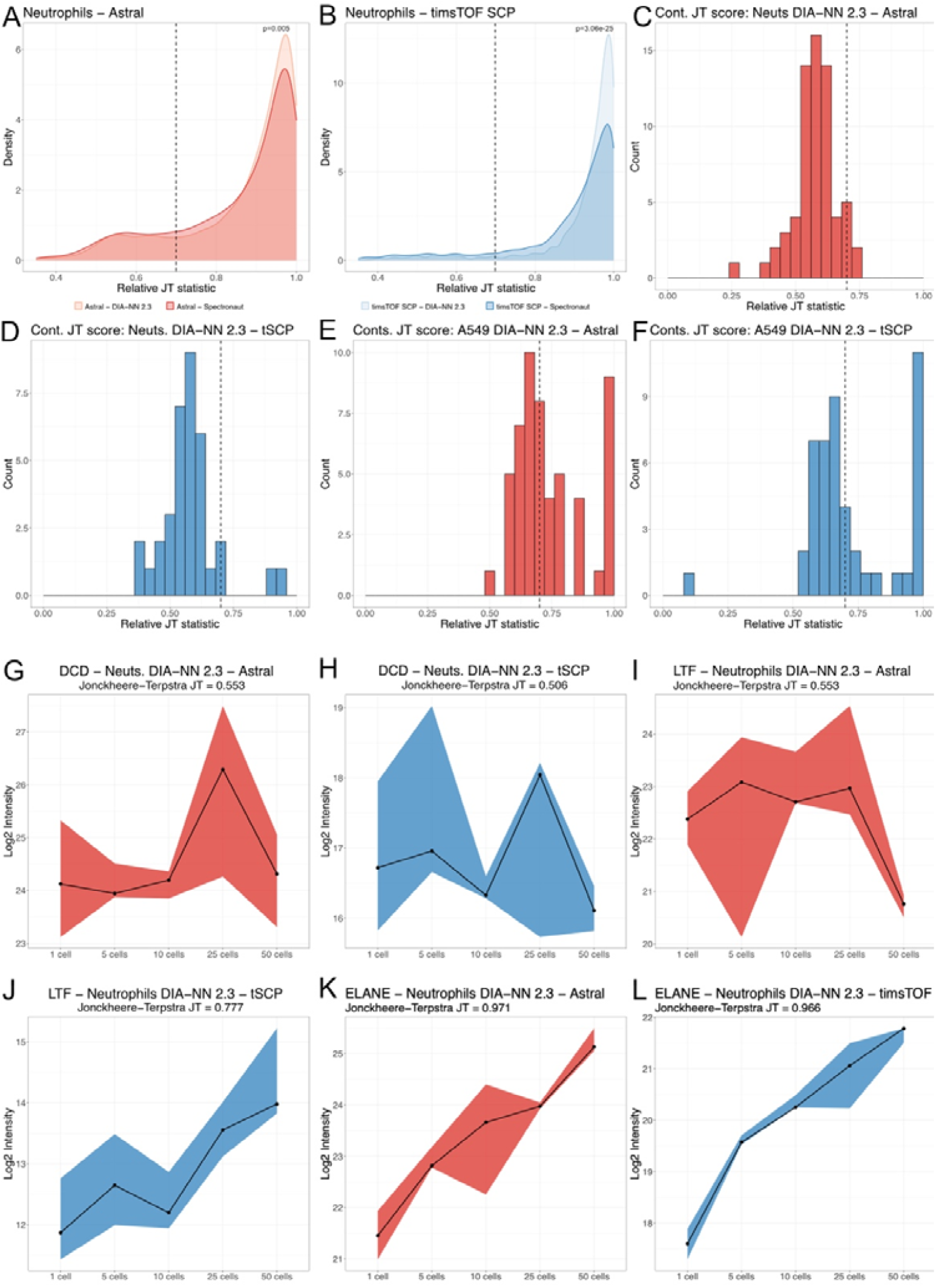
Potential skin contaminants affecting quantification in single cell proteomics: Density plots showing the relative Jonckheere-Terpstra (JT) score across Spectronaut and DIA-NN for the **(A)** Orbitral Astral (Astral) and **(B)** timsTOF SCP (tSCP). Histograms looking at the relative JT score contaminant proteins for DIA-NN outputs across **(C)** neutrophils on the Astral, **(D)** neutrophils on the tSCP, (E) A549 on the Astral and **(F)** A549 on the tSCP. Line plots showing the total intensity across the different number of neutrophils for **(G)** dermicidin (DCD) on the Astral **(H)** DCD on the tSCP, **(I)** lactoferrin (LTF) on the Astral, **(J)** LTF on the tSCP, **(K)** neutrophil elastase (ELANE) on the Astral and **(L)** ELANE on the tSCP. The dots represent the median intensity across the cell numbers and the ribbons the extent of the 95% confidence interval of the intensity.

Because many skin-associated proteins are not typically included in standard contaminant databases, we explored these results further. One clear example is dermcidin (DCD), an antimicrobial peptide found on the skin. In both the Astral and timsTOF SCP datasets, DCD showed a relative JT score below 0.56 (Fig. 8G&H), indicating no monotonic increase with increasing cell number. This behaviour is consistent with that expected for contaminant proteins. However, DCD is relatively easy to detect and filter because its abundance can be extremely high, even exceeding that of the most abundant neutrophil proteins in single-cell proteomics data. We therefore asked how many other proteins showed a similar pattern.

The tSCP data produced 1,920 proteins with valid relative JT scores and found that only 6.5% had a relative JT score <0.7 (Fig. 8A). The Astral data produced more proteins with valid relative JT scores, a total of 2,685, however a much higher proportion of these (16.5%) displayed contaminant like behaviour with a score <0.7 (Fig. 8B). Importantly, some these proteins are widely expressed and are therefore not included in the standard contaminant database. Examples include malic enzyme 1 (ME1) and insulin degrading enzyme (IDE), both of which are detected across a broad range of cell types. This category also includes neutrophil granules proteins like lactoferrin (LTF), which is known to be expressed by neutrophils^29,41-43^, but is also highly abundant in the skin^44^. When we examined the relative JT score for LTF in the Astral data, it showed contaminant-like behaviour with a score of 0.55 (Fig. 8I). The tSCP data looked more robust with a JT score of 0.77 (Fig. 8J), however it still clearly shows quantitative disruption and is far from the robust quantification seen from other granules like ELANE with relative JT scores higher than 0.97 (Fig. 8K&L).

### DIA-NN 2.5 and carafe improve the value of high-load libraries

The samples analysed in this study were searched using Spectronaut 20.3 and DIA-NN 2.3. However, a newer version of DIA-NN, reported to offer improved performance, was subsequently released. We therefore reanalysed the neutrophil samples using DIA-NN 2.5. Two searches were performed for both the Astral and tSCP datasets: one using only the single-cell data, and a second including both the single-cell data and all high-load libraries.

Our data revealed that DIA-NN 2.5 has limited improvements when searching only the single cell data. It show a small increase of 3.2% in the number of proteins identified on the Astral (Fig. 9A), while the tSCP data shows no significant differences between DIA-NN 2.3 and DIA-NN 2.5 (Fig. 9B). However, when we evaluated the results of the search which included all high-load libraries combined with the single cell data the results were very different. The data showed a 10.8% increase on the number of proteins identified on the Astral (Fig. 9C) while the CVs remained unchanged (Fig. 9D). On the tSCP the results showed an impressive increase of 32.5% protein identifications (Fig. 9E), while also maintaining equivalent CVs (Fig. 9F).

**Figure 9.**
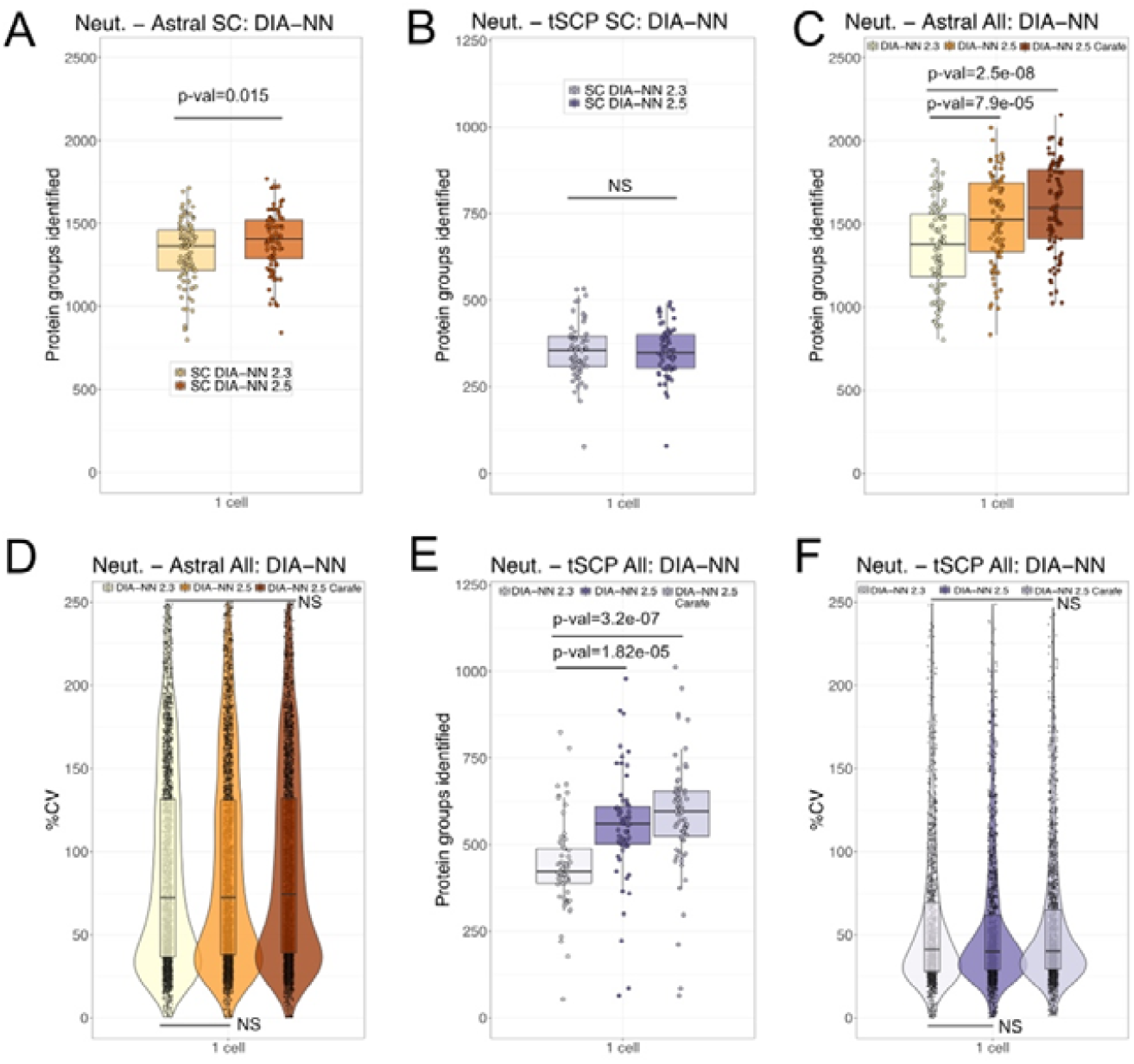
DIA-NN 2.5 and carafe improve the effect of high-load libraries: Boxplots showing the number of proteins identified in neutrophils when searching only single cell data on DIA-NN 2.3 and 2.5 for the (A) Orbitrap astral data (B) timstof SCP (tSCP) data. (C) Boxplot showing the number of proteins identified when the single cell and all high-load library samples are searched across the different DIA-NN versions for the Astral data. (D) Violin and box plots showing the coefficient of variation for all proteins across the different DIA-NN version for the Astral data. (E) Boxplot showing the number of proteins identified when the single cell and all libraries are searched across the different DIA-NN versions for the tSCP data. (F) Violin and box plots showing the coefficient of variation for all proteins across the different DIA-NN version for the tSCP data.

Finally, we tested the effect of Carafe^45^ on the search results from DIA-NN. Carafe is a tool that generates high-quality, experiment-specific in silico spectral libraries by training deep learning models directly on DIA data. We used a subset of raw files to generate the deep learning trained libraries for both the Astral and the tSCP with the same parameters the DIA-NN libraries had been generated. The data showed that using the carafe generated library with DIA-NN 2.5 improved protein IDs by an additional 4.4% on the Astral (Fig. 9C) while maintaining equivalent CVs (Fig. 9D). Similarly with the tSCP data it improved protein IDs by an additional 6.6% (Fig. 9E) while also maintaining comparable CVs (Fig. 9F)

Taken together, DIA-NN 2.5 seems to provide a modest boost when searching only single cell only. However, it provides very significant identification gains when adding high-load libraries both to the Astral and the tSCP. These performance gains are further improved by using a carafe generated library.

## Discussion

Single cell proteomics (SCP) has been revolutionised by the increased sensitivity of latest generation of instruments combined with improved sample processing methods. SCP is now rapidly expanding to provide insights into health and disease. Therefore, we set out to provide an overview relating to DIA search engines, software parameters, high-load libraries, quantitation and the effect of contaminants. Our work provides a comprehensive overview across cell types of different size and protein content, covering the range of most current single cell proteomic experiments and expands on recent benchmarking work done by others^46,47^.

Single cell work is used to define population or states, frequently by the presence or absence of proteins. This makes it incredibly important to ensure that the data is searched with robust search parameters that don’t wrongly propagate protein identifications^30^. In this work we highlight that the stringent single cell specific software parameters we previously used^25^, and included in the methods, produce only a modest reduction in IDs while maximising data quality and should be used for the vast majority of single cell experiments.

In this study, we compare the two most widely used instrumentation platforms for single-cell proteomics, the Astral and timsTOF, alongside two leading software tools, DIA-NN and Spectronaut, evaluating both protein identification rates and quantification precision. Our data confirms the higher sensitivity of the Astral compared to the timsTOF SCP (tSCP), which is particularly pronounced in smaller cells <60pg such as neutrophils. However, this sensitivity difference diminishes with increasing sample input, with the two platforms reaching comparable performance at approximately 3 ng of total protein injected. It is important to note that the tSCP is an older platform, and newer timsTOF instruments with improved sensitivity are now available.

Interestingly across both software tool we discovered that the tSCP produces much more precise data compared to the Astral. The most pronounced difference was detected in neutrophils, where the Astral data produced a median protein coefficient of variation (CV) >100%, where the tSCP was produced ∼70%, showing a very significant reduction compared to the Astral. This suggests the PASEF capabilities of the tSCP are particularly helpful for smaller cells.

The software comparison showed that Spectronaut 20.3 consistently produced higher protein identification rates than DIA-NN 2.3, in line with previous results^47^. This difference was most pronounced in the tSCP data when searches were performed using only single-cell runs. In contrast, DIA-NN produced significant improvements in CVs for the Astral data. Interestingly, the reduction in CVs was much less apparent in tSCP data, again suggesting that PASEF may reduce the additional precision gains provided by QuantUMS-based quantification. However, it is important to emphasise that CVs provide only a crude and sometimes incomplete measure of quantitative performance, as they can be artificially reduced while still producing poor-quality data^48^. A clear example of this is provided by the relative Jonckheere–Terpstra (JT) scores. Although DIA-NN produced little to no improvement in CVs for tSCP data compared with Spectronaut, a different pattern emerged when we assessed the proportion of proteins outside the quantitative range. Using a relative JT score threshold of <0.7 to define proteins whose quantification did not scale with cell number, only 6.5% of proteins detected with DIA-NN fell outside this range, compared with 11.7% for Spectronaut (Fig. 8B). Thus, despite near-identical CVs, the quantitative reliability of the detected proteins differed substantially between the two search engines.

Building on these software-dependent differences in quantitative performance, we also examined the effect of including high-load libraries in the search space. Rather than using diluted whole-cell lysates, these libraries were generated by sorting larger numbers of cells into separate samples, allowing us to assess their impact on both protein identifications and quantification. Interestingly, within Spectronaut, high-load libraries did not improve protein identification rates and, for tSCP data, libraries containing more than 20 cells actually reduced the number of protein identifications. In contrast, DIA-NN showed clear benefits from high-load libraries: they substantially increased protein identifications in the tSCP data and improved CVs for both tSCP and Astral datasets. This suggests that, when using DIA-NN, high-load libraries improve precision across both platforms, potentially by supplying improved training data for QuantUMS ^32^, thereby increasing the quantitative performance.

It is worth noting that results related to high-load libraries also depend on software parameters. The default setting in both DIA-NN and Spectronaut only enforce a 5% FDR across the different cells, enabling the transfer of potentially spurious identifications. The updated stringent parameters we propose limit this issue by enforcing a 1% FDR, but also reduce the value of the high-load libraries. Furthermore, the high-load library results have also changed considerably across the different software versions of DIA-NN and Spectronaut. There were considerable changes in Spectronaut when upgrading from version 19 to 20 and in DIA-NN from version 2.3 to 2.5. Specifically, Spectronaut 19 showed an underperforming FDR for single peptide proteins, which increased protein identification rates in general and increased the gains from using high-load libraries. This issue was rectified in Spectronaut 20, which in part explains the reduced value of high-load libraries for this version. Another important milestone has been the release of DIA-NN 2.5. In this DIA-NN version the performance of match-between-runs has been dramatically improved. This version produced only marginal gains when searching single-cell data without libraries, but substantially improved protein identification rates for both Astral and timsTOF SCP data when high-load libraries were included, even using the stringent search parameters, while maintaining comparable CVs.

We were also interested to assess how quantitative the identified proteins were and used a similar concept as the matched matrix background^37^, exploring the linear increases in the intensity of proteins when the number of cells increased. This led us to discover over 100 proteins seen in the neutrophil proteomes whose intensity did not scale well as the cell numbers increased, thus likely originating from contamination with the biggest source being skin. However, the challenge is that many of these proteins are exclusive to the skin proteomes and are shared with other cells as well, making them harder to identify and filter out.

This contamination can have pronounced quantitative effects in small cells <100pg of protein content, such as most immune cells, especially if the cells needed to be FACS sorted. The manual steps have the potential to increase the chance for skin contamination. It should be noted that this issue can be minimised, as in our previous work contaminant proteins like IL36G were barely detectable across single cells^25^. To assist the interpretation of the single cell data, we provide a database of these proteins and recommend careful interpretation of the results involving these. It should be noted that not all proteins will scale linearly if they reflect a functional state only seen in a small number of cells, hence we don’t fully recommend these being filtered out as full contaminants, but advise scrutiny. We also recommend adding cell dilution series to single cell experiment as we have done in this study. It adds few extra samples per plate, but it enables an analysis of how quantitative the detected proteins detected are.

Our work here provides guidelines on software, instrumentation platforms and high-load libraries across a wide range of cell types and cell sizes, and expect this to be of great use to the community when planning their single cell proteomics experiments.

## Methods

### Human participants

The collection of blood from healthy male and female control participants was approved by the Centre for Inflammation Research (CIR) Blood Resource Management Committee (AMREC #15-HV-013) and recruited from the University of Edinburgh CIR Blood Resource. Exclusion criteria for healthy controls included the following: infection with any blood borne diseases, previous or current intravenous drug abuse, anaemia, blood clotting disorders, anticoagulant drug therapy, regular use of steroids and/or under the age of 16 years old. Informed consent was obtained from all participants prior to sample collection. The study was approved by AMREC and the East of Scotland Research Ethics Committee REC and was conducted in accordance with the Declaration of Helsinki.

### Isolation of human blood neutrophils

Neutrophils were purified by dextran sedimentation and discontinuous plasma-Percoll gradients as described^49^.

### Fluorescence-activated cell sorting (FACS) of human blood neutrophils

From the Percoll gradients the purified neutrophils were sorted with FACS directly into 384 well plates to get the exact numbers required for proteomics. Following three washes with PBS w/o Ca2+ and Mg2+, single cells sorted as well as 5, 10, 25 and 50 neutrophils were sorted for proteomic analysis as high-load libraries. A FACS Aria Fusion sorter (Becton Dickinson) was used to collect cells in 384 well plates containing 1⍰µL of a cell lysis master mix consisting of 0.2% DDM (D4641-500MG, Sigma-Aldrich), 100⍰mM triethylammonium bicarbonate (TEAB; 17902-500ML, Fluka Analytical), 3⍰ng⍰µl^−1^ trypsin (Trypsin Gold, V5280, Promega) and 1% DMSO (83673.230, VWR Chemicals).

### Cell isolation and sample preparation of A549 cells

A549 cells were cultured at 37⍰°C in a humidified atmosphere at 5% CO_2_ using RPMI1640 medium supplemented with 10% fetal bovine serum (FBS; 10270106, Fisher Scientific), 1× penicillin–streptomycin (P0781-100ML, Sigma-Aldrich), 2mM GlutaMAX (35050061, Thermo Fisher Scientific) and 1⍰mM sodium pyruvate (11360070, Thermo Fisher Scientific). Cells were grown to around 75% confluency before trypsinization with 0.05% Trypsin-EDTA (25300-054, Thermo Fisher Scientific), followed by washing three times with PBS. Cells were resuspended in PBS at a density of 200 cells/μl for isolation with the cellenONE X1 Neo (Scienion).

A549 cell isolation, lysis and digestion were performed within a 384-well plate (Thermo Scientific Armadillo PCR Plate, 384 wells, 12657516) using the cellenONE X1 Neo robot as previously described^8^; using its control software (v3.2205.25.576API). Briefly, cells were sorted into wells containing 1⍰µl of master mix consisting of 0.2% DDM (D4641-500MG, Sigma-Aldrich), 1001mM triethylammonium bicarbonate (TEAB; 17902-500ML, Fluka Analytical), 3⍰ng⍰µl^−1^ trypsin (Trypsin Gold, V5280, Promega) and 1% DMSO (83673.230, VWR Chemicals). For single-cell samples, cells were deposited into individual wells, while for the 5-, 11-, 20- and 40-cell libraries, the respective cell number was sorted into a single well. Humidity and temperature were controlled at 50% and 12⍰°C during cell sorting. A549 cells were isolated with a diameter of 15–30⍰µm and 1.5 as maximum elongation. Cell lysis and protein digestion were performed at 501°C and 85% relative humidity for 30 min using adhesive tin foil to seal the plate before an additional 500⍰nl of 3⍰ng⍰μl^−1^ trypsin was added. The plate was incubated for another 1.5h and 2.5⍰μl of 0.1% TFA were added to the respective wells for quenching and storage at −70°C. For LC–MS/MS analysis, samples were directly injected from the 384-well plate after brief thawing and centrifugation.

### Vanquish Neo and Orbitrap Astral LC-MS/MS analysis

Samples were analyzed using the Thermo Scientific Vanquish Neo UHPLC system. Thermo Tune software version 0.4 or higher was used to acquire data. Peptides were separated on an Aurora Elite TS 15-cm nanoflow UHPLC column with an integrated emitter (Ion Optics) at 501°C using trap-and-elute mode with a PepMap (174500, Thermo Fisher Scientific). Peptide separation was performed at 50 samples-per-day (SPD). Fast sample loading was performed at a maximum pressure of 800 bar, using combined control and a maximum loading flow rate of 10µL/min. Solvent A was 0.1% formic acid (FA) in LC-MS grade water while solvent B was 80% acetonitrile, 0.1% formic acid in water with all being of LC-MS grade quality. The gradient flow started at 0 min with 0.45µL/min of 1% solvent B, at 0.1 min reaching 4% B, at 1.9 min reaching 12% B, at 2.0 min flow rate was changed to 0.2µL/min, at 12 min reaching 22.5% B, at 19.5 min reaching 40% B, at 22 min reaching 99% B and increasing the flow rate to 0.3µL/min and remaining at 99% B until the end of the run at 25 min.

For MS measuring, the Orbitrap Astral mass spectrometer Vanquish Neo and equipped with a FAIMS Pro Duo interface (Thermo Scientific) and an EASY-Spray source. 80°C and 100°C were used as outer and inner electrode temperature, respectively, as described previously to enhance peptide transmission^50^. A compensation voltage of –48⍰V was used for the FAIMS Pro Duo and an electrospray voltage of 1.9⍰kV for ionization.

On the Orbitrap Astral mass spectrometer, MS^1^ spectra were recorded for 27 min using the Orbitrap analyzer at a resolution of 240,000 from *m*/*z* 400 to 800 using an automated gate control (AGC) target of 500% and a maximum injection time of 100⍰ms. For MS^2^ in DIA mode using the Astral analyzer, nonoverlapping isolation windows of 20 *m*/*z* A scan range from an *m*/*z* of 400 to 800 was chosen. Precursor accumulation time was set to 60⍰ms and the AGC target to 800%.

### Cell isolation of RKO cells

Single cell seeding and processing was performed as previously described^51^. Briefly, 1000 nL of master mix (0.2% DDM (Sigma-Aldrich, D4641-5G), 100 mM TEAB (Supleco, 18597), 3 ng/µL trypsin (Thermo Scientific, 90058)) was dispensed into a 384 well plate (Eppendorf, 0030129547) inside the CellenONE. Cells matching the isolation criteria were dispensed to each well. Post-sorting the plate was sealed with a PCR foil seal (Thermo Fisher) and incubated at 50°C for 1.5 hours within a PCR cycler (Gene Amp, PE Appled Biosystems). Post incubation, the temperature was reduced to 20°C and 3.2 µL of 1 % formic acid (FA) was added to the samples.

### timsTOF SCP LC-MS/MS analysis

Samples were loaded onto purification and loading trap columns (Evosep, Evotips). Evotips were prepared according to manufacturer’s instructions. The samples were loaded onto the Evotips or manually from 384 well plate. After loading, the Evotips are washed with 0.1% FA followed by a final wash of 100 µL 0.1% FA and spinning for 10 seconds at 800 g. The samples are then transferred into Evosep One LC system for LC-MS/MS analyses.

LC–MS/MS analyses were performed using a timsTOF SCP mass spectrometer (Bruker) coupled to an Evosep One LC system (Evosep). Aurora Elite CSI analytical columns (IonOpticks; AUR3-15075C18-CSI) and a CaptiveSpray ionization source were used. Samples were analysed using the Whisper Zoom 40 SPD method, which employed a gradient flow of 200 nL/ min with a 31 min method duration. Eluted peptides were analysed with a parallel accumulation-serial fragmentation data-independent acquisition (diaPASEF) method. Method used for data acquisition consisted of 4 TIMS ramps with 5 mass ranges per ramp spanning from 327 to 1200 m/z and from 0.7 to 1.30 1/K0.

### Spectronaut search

With the parameters specified in Table X the raw files were searched on Spectronaut 20.3. Multiple searches were performed, all searched against a human SwissProt (September 2025) and a contaminants database. The single cell files were searched on their own, and then multiple searches including only 1 set of high-load libraries, i.e. either only single cells and the 5-cell libraries, or the single cell and the 25-cell libraries. Because of issues with Spectronaut it was ensured the high-load library raw files were always included first within the setup, as adding the single cell files first reduced protein identifications. One additional search was performed which included all single cells and all the high-load libraries for that search. This was done in the same manner across all 3 cell types.

### DIA-NN in-silico libraries

Two in-silico predicted libraries were created for the DIA-NN searches both using the parameters listed in Table 1. The first library was for the Orbitrap Astral data and limited precursor charge from 2-4, and precursor m/z to 400-800, reflecting the acquisition method for the instrument. A similar in-silico library was produced for the timsTOF SCP with precursor charge also 2-4 but precursor m/z from 350-1200. These libraries were used for all the DIA-NN searches.

**Table 1:** Spectronaut and DIA-NN parameters.

| DIA-NN | Values | Spectronaut Parameters | Values |
| --- | --- | --- | --- |
| Peptide length | 7-35 | Precursor Qvalue cutoff | 0.01 |
| Missed cleavages | 2 | Protein Qvalue cutoff (experiment) | 0.01 |
| Cross-run normalisation | Off | Protein Qvalue cutoff (run) | 0.01 |
| Quantification Strategy | QuantUMS (precision) | Precursos PEP cutoff | 0.15 |
| Proteotypicity | Genes | Protein PEP cutoff | 0.15 |
| Scoring | Generic | Peptide length | 7-35 |
| Machine learning | NNs (cross-validated) | Missed cleavages | 2 |
| FDR | 1% | Cross-run normalisation | off |
| Q.Value (parquet file) | 0.01 | Protein LFQ Method | MaxLFQ |
| PG.Q.Value (parquet file) | 0.01 | Major (protein) grouping | Protein Group ID |
| Lib.Q.Value (parquet file) | 0.01 | Minor (peptide) grouping | Stripped Sequence |
| Lig.PG.Q.Value (parquet file) | 0.01 | Major and Minor group top N | off |
| PG.PEP (parquet file) | 0.15 |  |  |
| MBR | enabled |  |  |

### DIA-NN search

As with Spectronaut the data were searched with DIA-NN 2.3, performing multiple searches for all three cell types. First the single cell were searched on their own, then individual searches adding the single cell files with only 1 high-load library at a time. This was repeated across all libraries (i.e. single cell and 5 cells, single cell and 10 cells…). For consistency library raw files were selected first before the single cell. For the Astral searches MS1 accuracy was set to 5ppm and MS2 accuracy to 10pm, for the timsTOF SCP searches MS1 accuracy to 15ppm and MS2 accuracy to 20ppm.

### Data analysis

All data analysis was carried out in R (v.4.5.0). The Spectronaut reports and the DIA-NN parquet files were used as starting input. The Spectronaut reports are already filtered to the required settings. The DIA-NN parquet files were filtered in R as follows: PG.Q.Value <=0.01, Q.value <=0.01, Lib.Q.Value<=0.01, Lib.PG.Q.Value<=0.01 and PG.PEP<=0.15.

### Outlier removal

The 5-cell high-load libraries RKO runs on the Orbitrap Astral show < 10 proteins identified and are removed from the analysis due to clear issues. One outlier in the neutrophil 25-cell libraries on the timsTOF SCP shows <500 proteins identified and is also removed as an outlier.

### Coefficient of Variation (CV)

All CVs were calculated as recommended in the community guidelines^31^. In short, the data without normalisation and without log transformation was fed into the proteomicsCV package version 0.4. The protCV() function was used on the non-log transformed non-normalised LFQ intensity.

### Jonckheere-Terpstra (JT) monotonicity score

A normalized Jonckheere–Terpstra (JT) score was used to quantify monotonic increases in total intensity across ordered cell-input groups. Intensities of NaN or 0 were filtered out. Only proteins detected in at least three high-load library groups containing at least two observed values were tested. All others were filtered out. The Jonckheere–Terpstra statistic was computed using the number of cells as an ordered factor and normalized by its maximum possible value for the observed group sizes.

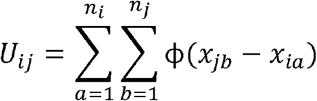

Where *x*_*ia*_ represents the intensity in group *i*, and *n*_*i*_ the number of non-missing values.*x*_*jb*_ represent the intensity in group *j*, and *n*_*j*_ the number of non-missing values.

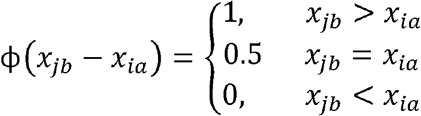

ϕ (*x*_*jb*_ − *x*_*ia*_) is the pairwise comparison function used to score the ordering between one observation from the higher-ordered group and one from the lower-ordered group.

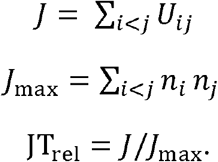

*J* is the overall Jonckheere-Terpstra statistic obtained by summing *U*_*ij*_ across all paired groups. *J*_max_ is the highest possible value of *J* given the group sizes and compositions. JT_rel_ normalises *J* providing a value between 0 and 1. A value of 0.5 denotes no increase in intensity when cell numbers increase, a value 1 of denotes a perfect increase in intensity as cell numbers increase and 0 a perfect decrease. JT_rel_<0.7 were considered poor.

### Significance testing

Significance testing for the coefficients of variation and the number of proteins identified were carried out in R using Welch’s T test which is two-tailed and doesn’t assume equal variance.

## Acknowledgments

Flow cytometry data were generated with support from the IRR Flow Cytometry and Cell Sorting Facility, within the University of Edinburgh. This work was supported by a Wellcome Early Career Award 327847/Z/25/Z (A.J.B), the infrastructure funding fourth call 2022/01 of the Austrian Research Promotion Agency (K.M.), the FWF projects 10.55776/PAT4142423 (K.M.), 10.55776/PAT4800425 (M.M.) and 10.55776/PAT2059025 (M.M.), the MRC MR/X01293X/1 (A.V.K) and BBSRC (BB/X019160/1) (A.V.K). We thank the Protein Chemistry Facility and acknowledge the VBCF for instrument access. For the purpose of open access, the author has applied a CC BY public copyright license to any Author Accepted manuscript version from this submission.

## Supplemental figures

**Supplemental figure 1.**
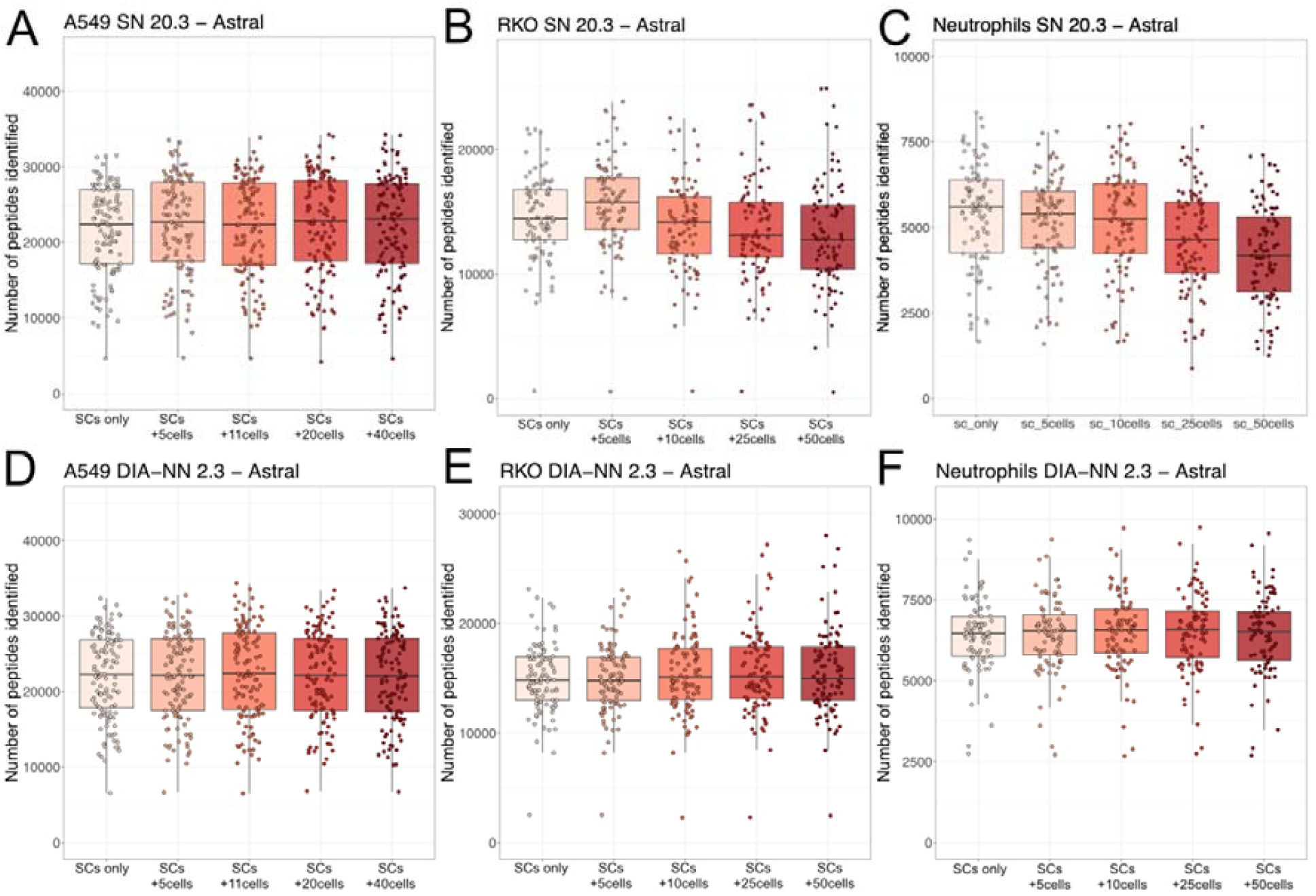
Astral peptide level IDs across high-load library search spaces: Boxplots showing the number of peptides identified on an Orbitrap Astral across the different search spaces which include single cells (SC) with the high-load libraries for **(A)** A549 cells with Spectronaut (SN), **(B)** RKO with SN, **(C)** neutrophils with SN **(D)** A549 cells with DIA-NN, **(E)** RKO cells with DIA-NN and **(F)** neutrophils with DIA-NN.

**Supplemental Figure 2.**
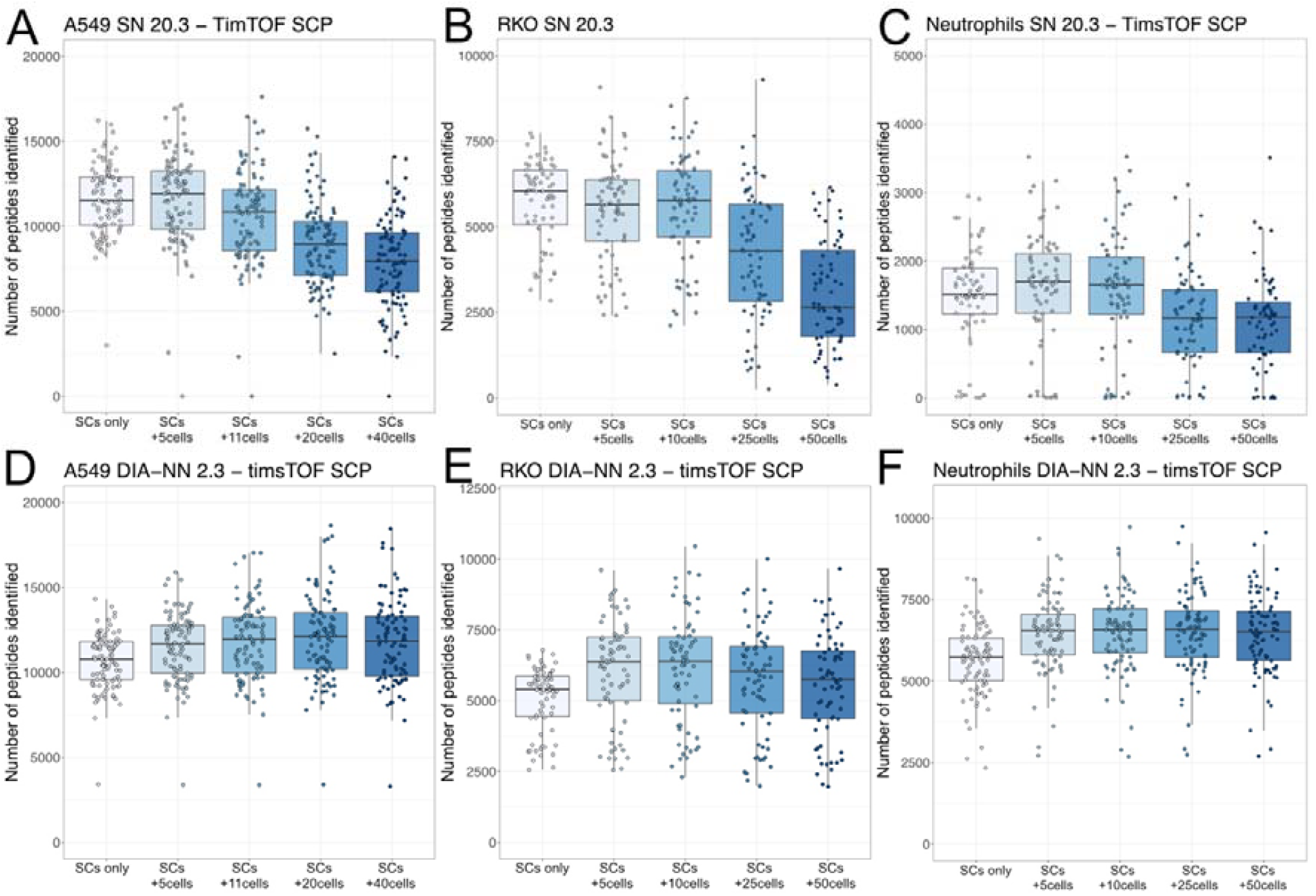
tSCP peptide level IDs across high-load library search spaces: Boxplots showing the number of peptides identified on a timsTOF SCP (tSCP) across the different search spaces which include single cells (SC) with specific high-load libraries for **(A)** A549 cells with Spectronaut, **(B)** RKO cells with Spectronaut, **(C)** neutrophils with Spectronaut, **(D)** A549 cells with DIA-NN, (E) RKO cells with DIA-NN and (F) neutrophils with DIA-NN

**Supplemental figure 3.**
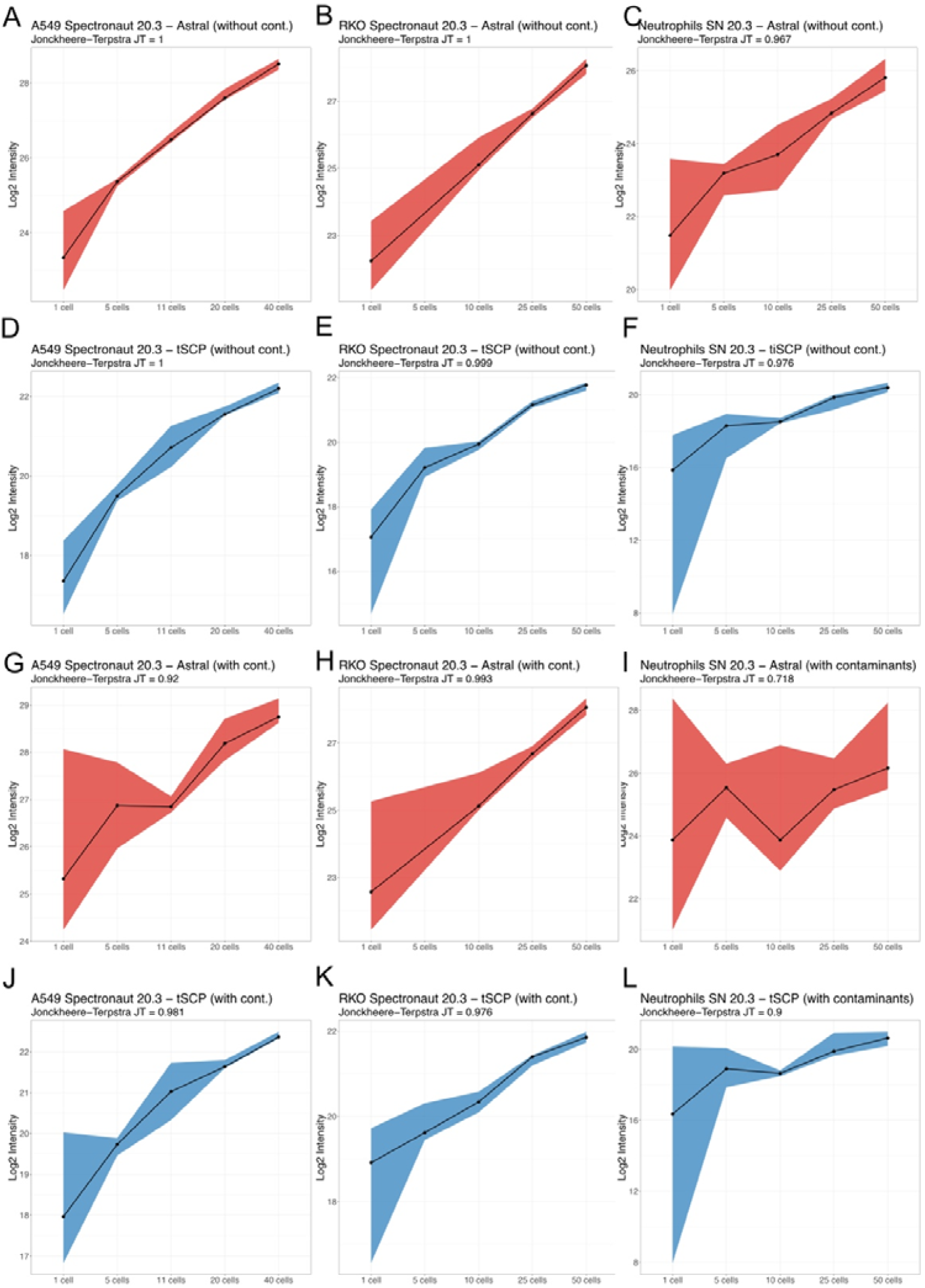
Spectronaut Intensity scaling with and without contaminants: Line plots showing the total intensity from Spectronaut across all proteins vs the different number of cells for **(A)** A549 on the Orbitrap Astral (Astral) without contaminants, **(B)** RKO on the Astral without contaminants, **(C)** Neutrophils on the Astral without contaminants, (D) A549 on the timsTOF SCP (tSCP) without contaminants, **(E)** RKO on the tSCP without contaminants, **(F)** neutrophils on the tSCP without contaminants, **(G)** A549 on the Astral with contaminants, **(H)** RKO on the Astral with contaminants, **(I)** Neutrophils on the Astral with contaminants, **(J)** A549 on the tSCP with contaminants, **(K)** RKO on the tSCP with contaminants, **(L)** neutrophils on the tSCP with contaminants. The dots represent the median intensity across the cell numbers and the ribbons the extent of the 95% confidence interval of the intensity.

## References

1 Rosenberger, F. A., Thielert, M. & Mann, M. Making single-cell proteomics biologically relevant. Nat Methods 20, 320–323 (2023). 10.1038/s41592-023-01771-9

2 Single-cell proteomics: challenges and prospects. Nat Methods 20, 317–318 (2023). 10.1038/s41592-023-01828-9

3 Budnik, B., Levy, E., Harmange, G. & Slavov, N. SCoPE-MS: mass spectrometry of single mammalian cells quantifies proteome heterogeneity during cell differentiation. Genome Biol 19, 161 (2018). 10.1186/s13059-018-1547-5

4 Specht, H. et al. Single-cell proteomic and transcriptomic analysis of macrophage heterogeneity using SCoPE2. Genome Biol 22, 50 (2021). 10.1186/s13059-021-02267-5

5 Brenes, A., Hukelmann, J., Bensaddek, D. & Lamond, A. I. Multibatch TMT Reveals False Positives, Batch Effects and Missing Values. Mol Cell Proteomics 18, 1967–1980 (2019). 10.1074/mcp.RA119.001472

6 Demichev, V., Messner, C. B., Vernardis, S. I., Lilley, K. S. & Ralser, M. DIANN: neural networks and interference correction enable deep proteome coverage in high throughput. Nat Methods 17, 41–44 (2020). 10.1038/s41592-019-0638-x

7 Bruderer, R. et al. Extending the limits of quantitative proteome profiling with data-independent acquisition and application to acetaminophen-treated three-dimensional liver microtissues. Mol Cell Proteomics 14, 1400–1410 (2015). 10.1074/mcp.M114.044305

8 Matzinger, M., Muller, E., Durnberger, G., Pichler, P. & Mechtler, K. Robust and Easy-to-Use One-Pot Workflow for Label-Free Single-Cell Proteomics. Anal Chem 95, 4435–4445 (2023). 10.1021/acs.analchem.2c05022

9 Ye, Z. et al. Enhanced sensitivity and scalability with a Chip-Tip workflow enables deep single-cell proteomics. Nat Methods 22, 499–509 (2025). 10.1038/s41592-024-02558-2

10 Ye, Z. et al. One-Tip enables comprehensive proteome coverage in minimal cells and single zygotes. Nat Commun 15, 2474 (2024). 10.1038/s41467-024-46777-9

11 Bubis, J. A. et al. Challenging the Astral mass analyzer to quantify up to 5,300 proteins per single cell at unseen accuracy to uncover cellular heterogeneity. Nat Methods 22, 510–519 (2025). 10.1038/s41592-024-02559-1

12 Petrosius, V. et al. Quantitative Label-Free Single-Cell Proteomics on the Orbitrap Astral MS. Mol Cell Proteomics 24, 100982 (2025). 10.1016/j.mcpro.2025.100982

13 Petrosius, V. et al. Exploration of cell state heterogeneity using single-cell proteomics through sensitivity-tailored data-independent acquisition. Nat Commun 14, 5910 (2023). 10.1038/s41467-023-41602-1

14 Ctortecka, C. et al. Automated single-cell proteomics providing sufficient proteome depth to study complex biology beyond cell type classifications. bioRxiv (2024). 10.1101/2024.01.20.576369

15 Ctortecka, C. et al. Automated single-cell proteomics providing sufficient proteome depth to study complex biology beyond cell type classifications. Nat Commun 15, 5707 (2024). 10.1038/s41467-024-49651-w

16 Woo, J. et al. High-throughput and high-efficiency sample preparation for single-cell proteomics using a nested nanowell chip. Nat Commun 12, 6246 (2021). 10.1038/s41467-021-26514-2

17 Zhu, Y. et al. Nanodroplet processing platform for deep and quantitative proteome profiling of 10-100 mammalian cells. Nat Commun 9, 882 (2018). 10.1038/s41467-018-03367-w

18 Leduc, A., Huffman, R. G., Cantlon, J., Khan, S. & Slavov, N. Exploring functional protein covariation across single cells using nPOP. Genome Biol 23, 261 (2022). 10.1186/s13059-022-02817-5

19 Ctortecka, C. et al. An Automated Nanowell-Array Workflow for Quantitative Multiplexed Single-Cell Proteomics Sample Preparation at High Sensitivity. Mol Cell Proteomics 22, 100665 (2023). 10.1016/j.mcpro.2023.100665

20 Demichev, V. et al. dia-PASEF data analysis using FragPipe and DIA-NN for deep proteomics of low sample amounts. Nat Commun 13, 3944 (2022). 10.1038/s41467-022-31492-0

21 Wu, T. et al. Single-cell proteomic landscape of the developing human brain. Nat Biotechnol (2026). 10.1038/s41587-025-02980-7

22 Fulcher, J. M. et al. Parallel measurement of transcriptomes and proteomes from same single cells using nanodroplet splitting. Nat Commun 15, 10614 (2024). 10.1038/s41467-024-54099-z

23 Sabatier, P. et al. Single-cell proteomics of pancreatic islet cells reveals type 1 diabetes and donor-specific features. bioRxiv (2025). 10.1101/2025.09.18.677038

24 Furtwangler, B. et al. Mapping early human blood cell differentiation using single-cell proteomics and transcriptomics. Science 390, eadr8785 (2025). 10.1126/science.adr8785

25 Sadiku, P. et al. Single cell proteomic analysis defines discrete neutrophil functional states in human glioblastoma. Nature Communications 17, 621 (2025). 10.1038/s41467-025-67367-3

26 Jiang, R. D., Shen, H. & Piao, Y. J. The morphometrical analysis on the ultrastructure of A549 cells. Rom J Morphol Embryol 51, 663–667 (2010).

27 Li, F., Cima, I., Vo, J. H., Tan, M. H. & Ohl, C. D. Single Cell Hydrodynamic Stretching and Microsieve Filtration Reveal Genetic, Phenotypic and Treatment-Related Links to Cellular Deformability. Micromachines (Basel) 11 (2020). 10.3390/mi11050486

28 Niemiec, M. J. et al. Trace element landscape of resting and activated human neutrophils on the sub-micrometer level. Metallomics 7, 996–1010 (2015). 10.1039/c4mt00346b

29 Long, M. B. et al. Extensive acute and sustained changes to neutrophil proteomes post-SARS-CoV-2 infection. Eur Respir J 63 (2024). 10.1183/13993003.00787-2023

30 Baker, C. P., Bruderer, R., Abbott, J., Arthur, J. S. C. & Brenes, A. J. Optimizing Spectronaut Search Parameters to Improve Data Quality with Minimal Proteome Coverage Reductions in DIA Analyses of Heterogeneous Samples. Journal of Proteome Research 23, 1926–1936 (2024). 10.1021/acs.jproteome.3c00671

31 Brenes, A. J. Calculating and Reporting Coefficients of Variation for DIA-Based Proteomics. J Proteome Res 23, 5274–5278 (2024). 10.1021/acs.jproteome.4c00461

32 Grossman, J. L. et al. Accurate quantification in proteomics with QuantUMS. Nat Biotechnol (2026). 10.1038/s41587-026-03131-2

33 Mahoney, D. W. et al. Relative quantification: characterization of bias, variability and fold changes in mass spectrometry data from iTRAQ-labeled peptides. J Proteome Res 10, 4325–4333 (2011). 10.1021/pr2001308

34 Karp, N. A. et al. Addressing accuracy and precision issues in iTRAQ quantitation. Mol Cell Proteomics 9, 1885–1897 (2010). 10.1074/mcp.M900628-MCP200.

35 (!!! INVALID CITATION !!! 6,14).

36 Krull, K. K., Ali, S. A. & Krijgsveld, J. Enhanced feature matching in single-cell proteomics characterizes IFN-gamma response and co-existence of cell states. Nat Commun 15, 8262 (2024). 10.1038/s41467-024-52605-x

37 Pino, L. K. et al. Matrix-Matched Calibration Curves for Assessing Analytical Figures of Merit in Quantitative Proteomics. J Proteome Res 19, 1147–1153 (2020). 10.1021/acs.jproteome.9b00666

38 Jin, H. et al. Systematic transcriptional analysis of human cell lines for gene expression landscape and tumor representation. Nat Commun 14, 5417 (2023). 10.1038/s41467-023-41132-w

39 Qi, L. et al. Integrated global proteomic and phosphoproteomic analysis of cisplatin-induced apoptosis in A549 cells. Biochem Biophys Res Commun 735, 150846 (2024). 10.1016/j.bbrc.2024.150846

40 Li, G. et al. Keratin gene signature expression drives epithelial-mesenchymal transition through enhanced TGF-beta signaling pathway activation and correlates with adverse prognosis in lung adenocarcinoma. Heliyon 10, e24549 (2024). 10.1016/j.heliyon.2024.e24549

41 Sollberger, G., Brenes, A. J., Warner, J., Arthur, J. S. C. & Howden, A. J. M. Quantitative proteomics reveals tissue-specific, infection-induced and species-specific neutrophil protein signatures. Sci Rep 14, 5966 (2024). 10.1038/s41598-024-56163-6

42 Hoogendijk, A. J. et al. Dynamic Transcriptome-Proteome Correlation Networks Reveal Human Myeloid Differentiation and Neutrophil-Specific Programming. Cell Rep 29, 2505–2519 e2504 (2019). 10.1016/j.celrep.2019.10.082

43 Reyes, L. et al./person-group> A type I IFN, prothrombotic hyperinflammatory neutrophil signature is distinct for COVID-19 ARDS. Wellcome Open Res 6, 38 (2021). 10.12688/wellcomeopenres.16584.2

44 Cumberbatch, M. et al. Regulation of epidermal Langerhans cell migration by lactoferrin. Immunology 100, 21–28 (2000). 10.1046/j.1365-2567.2000.00014.x

45 Wen, B. et al. Carafe enables high quality in silico spectral library generation for data-independent acquisition proteomics. Nat Commun 16, 9815 (2025). 10.1038/s41467-025-64928-4

46 Yang, Z., Ma, T. P., Choi, M., Yu, F. & Zhu, Y. Systematic Evaluation of Data-independent Acquisition Workflows for High-Throughput and Low-Input Proteomics Analysis with an Astral Mass Spectrometer. J Proteome Res 25, 1686–1699 (2026). 10.1021/acs.jproteome.5c01028

47 Wang, J. et al. Benchmarking informatics workflows for data-independent acquisition single-cell proteomics. Nat Commun 16, 10276 (2025). 10.1038/s41467-025-65174-4

48 Ivanov, M. V., Garibova, L. A., Postoenko, V. I., Levitsky, L. I. & Gorshkov, M. V. On the excessive use of coefficient of variation as a metric of quantitation quality in proteomics. Proteomics 24, e2300090 (2024). 10.1002/pmic.202300090

49 Haslett, C., Guthrie, L. A., Kopaniak, M. M., Johnston, R. B., Jr. & Henson, P. M. Modulation of multiple neutrophil functions by preparative methods or trace concentrations of bacterial lipopolysaccharide. Am J Pathol 119, 101–110 (1985).

50 Hoch, D. G., Belford, M., Heil, L. R., Mechtler, K. & Matzinger, M. Low-resolution FAIMS for increased peptide coverage in low-load and single-cell proteomics. Sci Rep 16 (2026). 10.1038/s41598-026-45228-3

51 Makar, A. N., Holkham, J., Lilla, S., Wilkinson, S. & von Kriegsheim, A. Overcoming Preservation Challenges to Enable Single-Cell Proteomics of Fixed Cells and Tissue Samples with Retained Proteome Integrity. J Proteome Res 24, 3666–3682 (2025). 10.1021/acs.jproteome.5c00268

